# Dual NMDAR signaling in astrocytes: flux-independent pH sensor & flux-dependent mitochondrial regulator through membrane-mitochondria communication

**DOI:** 10.1101/633875

**Authors:** Pavel Montes Oca Balderas, Arturo Hernández-Cruz

## Abstract

Glutamate N-methyl-D-aspartate (NMDA) receptor (NMDAR) is critical for neurotransmission as a Ca2+ channel. Nonetheless, several reports have also demonstrated flux-independent signaling. Astrocytes express NMDAR distinct from its neuronal counterpart, but cultured astrocytes have no electrophysiological response and controversial findings have questioned NMDAR function. We recently demonstrated that in cultured astrocytes NMDA at pH6 (NMDA/pH&) elicits flux-independent Ca2+ release from the Endoplasmic Reticulum (ER) and depletes mitochondrial membrane potential (mΔψ). Here we show that flux-independent Ca2+ release is mainly due to pH6, whereas mΔψ depletion requires both pH6 and flux-dependent NMDAR signaling. Immunofluorescence exhibited that plasma membrane (PM) NMDAR is apposed to ER and mitochondria or surrounds these organelles. Moreover, NMDA/pH6 treatment generated ER stress, increased endocytosis, mitochondria-ER and -nuclear contacts and strikingly, PM invaginations near mitochondria along with electrodense structures referred here as PM-mitochondrial bridges (PM-m-br). These data and earlier observations strongly suggest PM-mitochondria communication. As a proof of concept of this notion, NMDA/pH6 provoked mitochondria labeling by the PM dye FM-4-64FX. Finally, we analyzed by WB NMDAR subunit GluN1 to explore putative causes of NMDAR dual function, we found fragments with M.W. consistent with previously identified cleavage sites. Accordingly, GluN1 intracellular and extracellular domains presented little colocalization. Our findings demonstrate that NMDAR plays a dual function: a flux-independent pH sensor and a flux-dependent regulator of mΔψ. More importantly, mΔψ depletion seems to be mediated by PM-mitochondria communication. Finally, we found different GluN1 fragments that could be involved in NMDAR dual signaling, although causality awaits demonstration.

## INTRODUCTION

The glutamate (Glu) N-methyl-D-aspartate (NMDA) receptor (NMDAR) is traditionally conceived as an ionotropic receptor with a critical role in neurotransmission. This function requires co-agonist (Gly or D-Ser) binding for channel opening and is regulated by ions (Mg2+, Zn2+, H+), or other molecules (1–3). The NMDAR is a tetramer of homodimers or a heterotrimer conformed by two obligate subunits GluN1 coupled to GluN2 and/or GluN3 subunits. GluN1 is present in all NMDAR described so far, as it plays a central role for NMDAR assembly at the Endoplasmic Reticulum (ER) and its intracellular (IC) trafficking (1–3). However, NMDAR signaling is more complex, because Ca2+ inflow can switch on or off IC pathways with pro-survival or pro-death effects, suggested to be the result of synaptic or extra-synaptic NMDAR function, respectively (4). Moreover, several reports have also demonstrated flux-independent actions of this receptor (also termed metabotropic-like or non-canonical functions), that mediate long term depression (LTD) and other cellular events, through molecular mechanisms that have been only slightly investigated (5).

In astrocytes, NMDAR expression and function were matter of debate for several years. However, nowadays it is clear that astrocytes *in situ* express functional NMDAR, although it displays several singularities, given in part by its subunit composition (6, 7). Nevertheless, classical electrophysiological experiments found no ionic flux in cultured astrocytes, and controversial findings have questioned NMDAR function (6, 8). Nonetheless, we and others have recently demonstrated that the NMDAR can initiate flux-independent activities (9–11). We found Ca2+ release from the Endoplasmic Reticulum (ER) in response to NMDA 1mM at pH≈6 (NMDA/pH6). These levels of acidity and glutamate can be reached in the synaptic cleft after intense synaptic transmission. Indeed, more acidic pH values can be achieved in pathological conditions such as stroke, inflammation or hypercapnia (12–15). We demonstrated that iCa2+ rise was sensitive to competitive NMDAR inhibitors or GluN1 subunit Knock Down, demonstrating NMDAR involvement. Surprisingly, iCa2+ rise was insensitive to NMDAR pore blocker MK-801 or extracellular Ca2+ depletion, but it was blocked by InsP_3_R and RyR inhibitors. Also, NMDA/pH6 depleted mitochondrial membrane potential (mΔψ) (9). In this work, we further study the signaling, cellular biology and biochemistry of the NMDAR to gain insight into its complex biology, that given its expression and diversity in different cells and tissues (16), it seems to be greater than conceived. We show here that NMDAR has a dual signaling function as a flux-independent pH sensor or as a flux-dependent channel that together with acid pH depletes mΔψ. In addition, our experiments along previous reports, strongly suggest that mΔψ is achieved by plasma membrane (PM) -mitochondria communication through structures referred here as PM-mitochondrial bridges (PM-m-br). The communication between these organelles has been studied in yeast but to our knowledge only one study has reported it in mammalian cells (17, 18). Finally, in an effort to explore the biochemistry behind these findings, we found GluN1 fragments that could be involved in the dual signaling function of the NMDAR, although causality awaits demonstration.

## METHODS

### Rat Cultured Cortical Astrocytes (rCCA)

rCCA were prepared as described before (19) with modifications. Wistar rats (*Rattus norvegicus*) were decapitated on postnatal day 0-3 accordingly to guidelines established by the Institutional Committee for Laboratory Animal Care and Use from the National Institute for Neurology and Neurosurgery that approved this study (Project 94/12). Cortices were dissected in cold Hank’s Balanced Salt Solution (HBSS) and dissociated. The cells were recovered, washed and seeded (2.5×10^4^ cells/cm^2^) in culture medium (DMEM with 10% FBS, 100 U/ml penicillin and 100 ng/ml streptomycin) onto poly-D-lysine (PDL) covered tissue culture flasks. The medium was replaced after 24 h and every three days thereafter. At confluence (12-14 days), flasks were shaken to detach microglia and astrocytes were harvested with trypsin and seeded at 8×10^3^ cells/cm^2^ in appropriate dishes or coverslips for the indicated experiment. The cells were not passed more than 3 times and were used for the indicated experiments 2-8 days after seeding.

### Reagents and antibodies

Salts, reagents, NMDA, D(−)-2-amino-5-phosphonopentanoic acid (APV), Kynurenic acid (KYNA), Dizocilpine (MK-801) and cell culture media powder were from Sigma Chemical Co. (St. Louis, MO, USA). Media supplements, Hoechst, 5,5’,6,6’-tetrachloro-1,1’,3,3’-tetraethylbenzimidazolylcarbocyanine iodide (JC-1), Fluo-4 acetomethylester (Fluo-4-AM), Mitotracker Deep Red FX (MTR), ER- Tracker Red (ERT), FM 4-64FX and Dextran labeled with Alexa Fluor 647 were from Life Technologies (Carlsbad, CA, USA). Culture plates were from Costar (Tewksbury, MA, USA). Antibodies (Abs) against NMDAR subunits were purchased from Santa Cruz Biotechnology (Santa Cruz, CA, USA). Secondary Abs were from Jackson Immunochemicals (West Grove, PA, USA). Western Blot (WB) reagents were from Bio-Rad (Hercules, CA, USA), Enhanced Chemiluminiscent (ECL) reagents were from Thermo Scientific (Rockford, IL, USA), and Hybond™ and Hyperfilm™ were from GE Healthcare Amersham (Piscataway, NJ, USA). Protease Inhibitory cocktail was from Roche (Basel, CH).

### Intracellular calcium (iCa2+) measurements

iCa^2+^ determinations were performed as described previously (9) with modifications. rCCA seeded onto coverslips were loaded for 45 min with 1 μM Fluo-4-AM/0.02% pluronic acid in Hank’s Balanced Salt Solution (HBSS) containing 137 mM NaCl; 3mM KCl; 336 μM Na2HPO4; 442 μM KH2PO4; 5.55 mM Glucose; 1.2 mM CaCl_2_, 400 μM MgSO_4_ and 440 μM MgCl_2;_ pH 7.3-7.4. Coverslips were washed 3X and transferred to a stage chamber and fluorescence emission was recorded with a 10X 0.4 N.A. objective from an Olympus IX-81 (Olympus Corporation, Japan) microscope equipped with a Hamamatsu Orca Flash 2.8, running with Cell Sens Dimension software (Olympus Corporation, Japan) and mercury lamp illumination. Fluorescence was recorded with a 485/30 bandpass excitation filter, a 505 long-pass dichroic mirror and a 510/50 bandpass emission filter. Time-lapse recordings were acquired at 12 frames per min (f/m)(0.2 Hz). After 120 sec recording basal fluorescence with HBSS perfusion at 900 μl/min applied with a syringe injector (New Era Pump Systems Inc; NY, USA), with or without inhibitors, NMDA or HBSS at the indicated pH was perfused and recording continued for 230 sec, to finally return to vehicle perfusion for 30 sec. Before the experiment, cells were treated 15-45 min with inhibitors that were also constantly perfused during the recording. The perfusate was applied with a fixed 0.8 mm needle placed 700 μm above the field of view. Time–lapses were analyzed with Cell Sens Dimension software; background noise was subtracted and oval regions of interest (ROI) were drawn around each cell soma. Cells showing bright cytoplasmic puncta were discarded. The mean fluorescence intensity (F) was obtained for each ROI and iCa^2+^ responses were evaluated by calculating the relative fluorescence change (ΔF/F_0_) with the formula ΔF/F_0_ = (F*t*_*x*_−F_0_)/F_0_, where F*t*_*x*_ is F at time *x* and F_0_ is the average fluorescence in frames 5-10. Data are presented as average response traces with cells obtained from at least three different cultures. The integrated relative fluorescence change ∫(ΔF/F_0_) for each cell was calculated as the sum of ΔF/F_0_ values between sec. 50-350. The average ∫(ΔF/F_0_) values, number of cells (*n*), experiments and numerical descriptors for averaged response, are found in Table I. ∫(ΔF/F_0_) values for individual cells are presented as raster plots and distribution histograms for each experiment in supplementary Figure 1.

**TABLE I.** Data summaries from iCa2+ averaged responses Fig. 1.

|  | **NMDA/pH7** | **NMDA/pH6** | **NMDA/pH5** | **HBSS/pH7** | **HBSS/pH6** | **HBSS/pH5** | **APV** | **KYNA** |
| --- | --- | --- | --- | --- | --- | --- | --- | --- |
| **# of cells (n)/**<br>**# of**<br>**experiments** | 130/4 | 136/4 | 335/7 | 210/4 | 335/7 | 495/9 | 249/3 | 551/11 |
| **Average**<br>**($\int(\Delta F/F_0) \pm \text{SEM}$)** | 2.81±0.35 | 5.5±0.45 | 3.37±0.29 | 0.62±0.28 | 5.86±0.3 | 1.73±0.16 | 1.29±0.18 | 1.21±0.16 |
| **% response to**<br>**normalized**<br>**NMDA/pH6** | 51.1 | 100 | 61.3 | 11.3 | 106.5 | 31.4 | 23.38 | 21.91 |
| ***p* value**<br>**compared with**<br>**NMDA/pH6** | 3.65X10<sup>-6</sup> | N.A. | 8.22X10<sup>-5</sup> | 2.72X10<sup>-17</sup> | 0.47 | 5.14X10<sup>-21</sup> | 8.9X10<sup>-22</sup> | 6.38X10<sup>-26</sup> |

**Figure 1.**
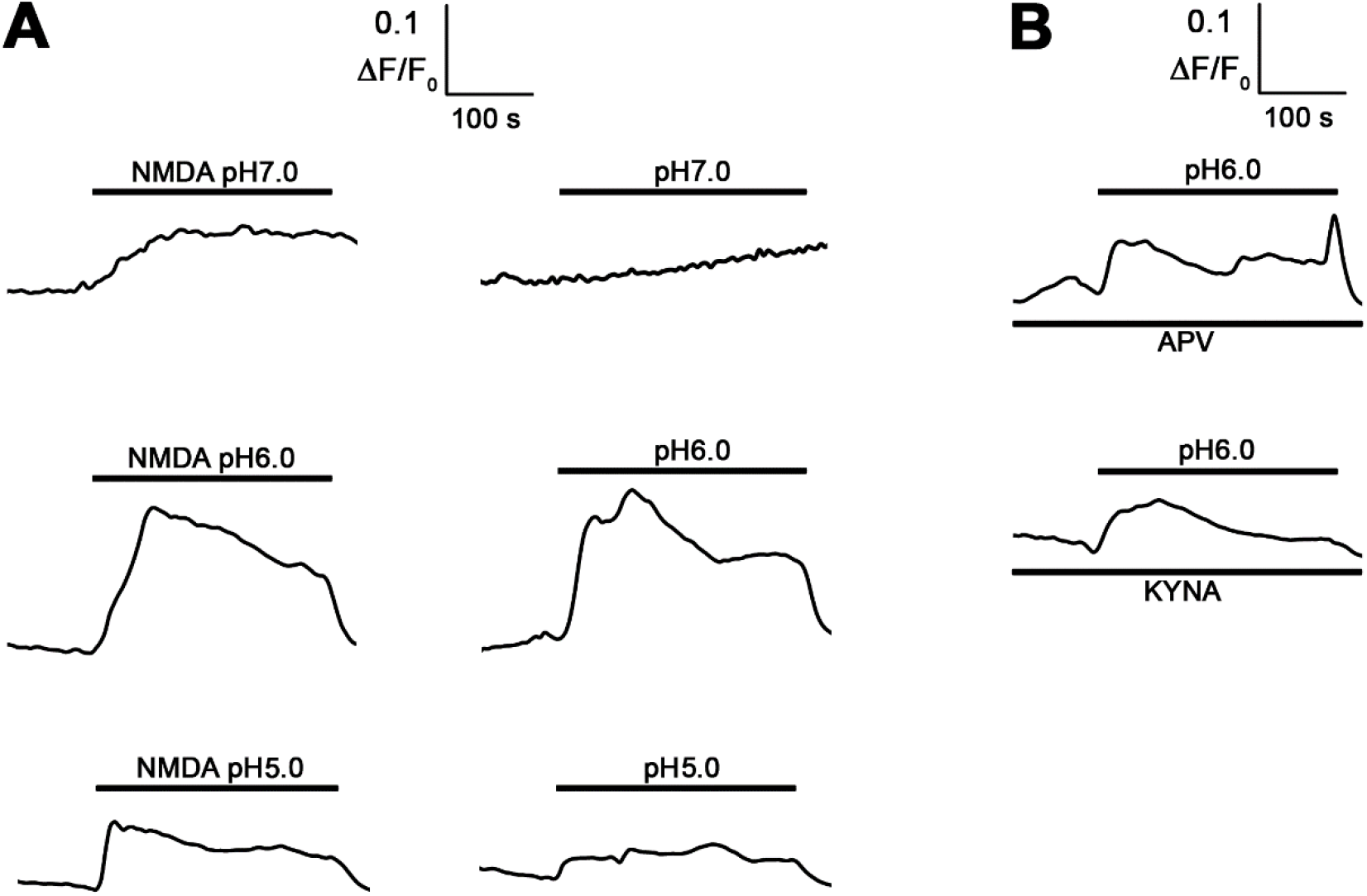
Effect of pH on iCa2+ rise of rCCA elicited by NMDA/pH6. **A** *left column*. ΔF/F_0_ averaged response of Fluo-4 labeled rCCA perfused with NMDA at different pH values as indicated. **A** *right column*. ΔF/F_0_ averaged response of Fluo-4 labeled rCCA perfused with HBSS at different pH values as indicated. **B** ΔF/F_0_ averaged response of Fluo-4 labeled rCCA perfused with HBSS/pH6 in presence of APV and KYNA, NMDAR competitive inhibitors of the Glu and Gly sites, respectively. Bars above each curve indicates treatment and its duration. Bars below indicate the inhibitor tested.

### Mitochondrial Membrane Potential (mΔψ) monitoring

rCCA seeded onto 12 mm coverslips were labeled with 600 ng/ml of JC-1 (1H-Benzimidazolium, 5,6-dichloro-2-[3-(5,6-dichloro-1,3-diethyl-1,3-dihydro-2H-benzimidazol-2-ylidene)-1-propenyl]-1,3-diethyl-iodide) for 15 min at 37°C in culture medium. Cells were washed 3X with HBSS and placed in the bottom of a plexiglass recording chamber attached to the stage of a custom-made spinning disc confocal microscope (Solamere Technology Group, Salt Lake city, USA) coupled to an upright microscope (Nikon Eclipse 80i; Nikon Corp., Tokyo, Japan) and continuously perfused (1 ml/min) with HBSS applied to the recording chamber with a syringe injector. JC-1 was excited at 488nm and 561nm with monochromatic light from solid state Coherent Obis lasers (Laser Physics, Reliant 100 s488, West Jordan, UT), coupled to a Yokogawa spin-disk confocal scan head (CSU10B, Yokogawa Electronic Co., Tokyo, Japan and Solamere Technology Group, Salt Lake city, USA). Emitted fluorescence was band-passed (525/50, 605/52) before collection with a cooled digital CCD camera (Andor Technology iXon 897, Oxford Instruments, High Wycombe, UK). The full system was controlled with the open source microscopy software Micromanager (20). Time-lapse recordings were acquired at 0.5 Hz for 180 sec. After obtaining basal readings for 60 sec in HBSS with or without inhibitors, cells were treated with 1 mM NMDA or HBSS at the indicated pH and recorded for 2 additional min. The solutes were applied directly to the field of view with a gravity-fed (≈400 μl/min), homemade micro-perfusion system consisting of a 0.8 mm syringe tip placed 500 μm above and at the edge of the field of view with an Eppendorff micromanipulator (Eppendorff, Germany). Time—lapse recordings were analyzed with Cell Sens Dimension software (Olympus Corporation, Japan). Background noise was subtracted and ROIs were drawn around each cell soma. The average fluorescence intensity for green (*x*F_*i*_g) and orange-red (*x*F_*i*_or) channels were obtained for each ROI and mΔψ was calculated as ratios [mΔψ=(*x*F_*i*_or)/(*x*F_*i*_g)] (subsequently referred to as mΔψ). Data are presented as average mΔψ. The diversity of individual cellular responses is presented as raster plots for each experiment in supplementary Figure 2. In addition, distribution histograms were constructed with the rate between t180/t0 for each cell (Supplementary Figure 2).

### ER and mitochondria labeling

ER and mitochondria were labeled with ERT (ER- Tracker Red) or MTR (Mitotracker Deep Red FX), respectively, according to manufacturer instructions. Briefly, rCCA seeded onto coverslips were washed 3X with HBSS and incubated for 30 min with 1 μM ERT or 5 μM MTR in HBSS at 37°C. Then, cells were washed 3X and subject to immunofluorescence as described below. *z*-stacks of MTR labeled cells were acquired with a 100X/N.A. 1.3 objective mounted on the spinning disc microscope described above using an excitation Laser of 640 nm and an emission filter 700/75 nm. *z*-stacks of ERT labeled cells were acquired with a 60X/N.A. 1.3 objective in a DMI 6000 inverted microscope (Leica Microsystems, Wetzlar, Germany) equipped with a Leica EL6000 external mercury light source connected via a liquid light guide coupled to a CCD (DFC345 FX; Leica Microsystems), with filter cube Y5 from Leica (Ex: 620/60; Em:700/75; DM:660), and LE software. For image analysis, background was subtracted and colocalization was inspected visually (Olympus Corporation, Japan).

### Immunofluorescence

Cells seeded onto coverslips were washed with PBS, fixed with PBS-4% paraformaldehyde (PFA)-4% sucrose on ice and then permeabilized with PBS-0.5% Tween-20. For some experiments the NMDAR present in the plasma membrane was detected skipping the permeabilization step. After washing, cells were incubated with primary Ab (1 μg/ml) one hour in M1 buffer (140 mM NaCl, 20 mM HEPES, 1 mM Cl_2_Ca, 1 mM Cl_2_Mg, 5 mM KCl), washed and then incubated with secondary Ab (46.6 ng/ml) one hour. Nuclei were labeled with Hoechst 33342 (10 μM). Coverslips were washed and mounted with homemade Mowiol-DABCO. *z*-stacks of double immunofluorescence experiments were acquired (60X/N.A. 1.35 objective) in the Olympus microscope described above equipped with motorized stage, that was corrected for z-shift between different wavelengths with fluorescent beads of 100 nm. For image analysis, background was subtracted and colocalization was visually inspected.

### Electron Microscopy (EM)

Cells seeded onto 10 cm petri dishes were washed with HBSS and incubated 30 min with HBSS/pH6 or NMDA/pH6. Cells were then washed 3X and fixed with PBS-2.5% glutaraldehyde for 2h at 4°C. Then cells were harvested and washed 3X with ice cold PBS 15 min and post-fixed with 1% OsO4 for 1 h at 4°C. Cells were washed and dehydrated with increasing concentrations of cold ethanol. Cell samples were incubated with acetonitrile 2X for 15 min and then with acetonitrile-Epon for 48 h in a desiccator. The samples were embedded in fresh Epon and polymerized in a stove for 48 h at 60°C. Samples were cut at 70-80 nm with a Reichert Ultracut ultramicrotome (Leica Microsystems, Wetzlar, Germany) and contrasted with 2% uranyl acetate for 20 min and lead citrate 2% for 10 min. Samples were observed in a Transmission Electron Microscope JEOL JEM 1200 EX II (Peabody, Massachusetts, USA).

### Endocytosis Assays

Cells seeded onto coverslips were washed with HBSS 3X and then incubated 15 min with 500 μg/ml of Dextran 10,000 labeled with Alexa Fluor 647 with or without NMDA/pH6. Then, cells were washed 3X with ice cold PBS and fixed 10 min with ice cold PBS-4% PFA-4% sucrose. *z*-stacks were acquired with the IX-81 Olympus microscope set up described above with a 60X/N.A. 1.35 objective, an excitation band pass filter (640/30), a 660 nm dichroic mirror and a band pass emission filter (700/75). For image analysis, background was substracted and z-stacks were processed with software deconvolution algorithm, then the channel signal was segmented and detected particles were counted in ROIs drawn around cell soma with Cell Sens software (Olympus Corporation, Japan) in the slice focused at the basal membrane.

### FM 4-64FX labeling

To analyze FM 4-64 labeling of mitochondria, rCCA prelabeled with MTR as indicated above were incubated for 10 min with FM 4-64FX (5 μg/ml) with or without NMDA/pH6 while recording with the Spinning Disc microscope described above with a 488 Laser for excitation and an emission filter of 700/75 nm. Simultaneously, MTR was recorded as described above. For image analysis, background was subtracted and colocalization was inspected visually with Cell Sens software (Olympus Corporation, Japan).

### Western Blot

Cells were washed with cold PBS, lysed with reducing sample buffer containing protease inhibitors and boiled 5 min. Lysates were resolved on SDS-polyacrylamide gels, and proteins were transferred to a nitrocellulose membrane. Protein detection was conducted by incubating with primary Ab (66 ng/ml) one hour, then with the appropriate secondary Ab (26 ng/ml) one hour and then revealed by ECL (Thermo Fisher Scientific, USA). The blot was stripped with Stripping Buffer (Thermo Fisher Scientific, USA) and re-probed after detection controls were performed.

### Statistical analysis

Statistical *p* values for iCa^2+^, mΔψ and endocytosis experiments were determined by student’s t-test with Excel (Microsoft). In agreement with recent analysis of statistical *p* value use that may mislead data interpretation (21), no threshold is assigned to claim statistical significance. *p* values for iCa2+ and mΔψ experiments are shown in Table I and Table II, respectively, or in Figure 6 for endocytosis assays.

**TABLE II.** Data summaries from mΔψ averaged responses in Fig. 2.

|  | **NMDA/pH6** | **HBSS/pH6** | **NMDA/pH7.3** | **APV 10 μM** | **APV 100 μM** | **MK-801** |
| --- | --- | --- | --- | --- | --- | --- |
| **# of cells (n)/**<br>**# of**<br>**experiments** | 326/4 | 192/4 | 238/7 | 89/9 | 298/3 | 123/11 |
| **Ratio**<br>**t180/t0±SEM** | 0.83±0.02 | 0.98±0.02 | 1.06±0.01 | 0.92±0.02 | 0.98±0.02 | 1.02±0.01 |
| ***p* value**<br>**compared with**<br>**NMDA/pH6** | N.A. | 2.99X10<sup>-6</sup> | 2.54x10<sup>-13</sup> | 0.046 | 2.3x10<sup>-6</sup> | 1.89x10<sup>-6</sup> |

**Figure 2.**
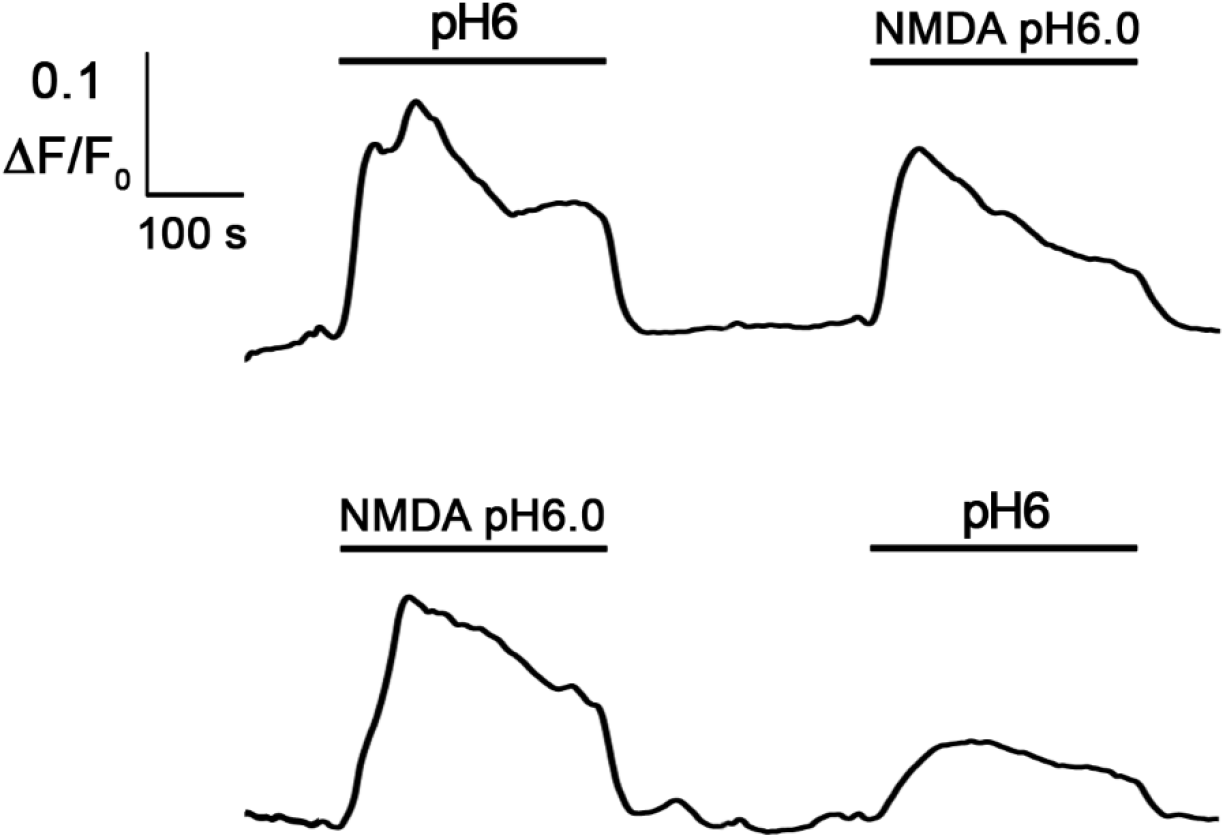
NMDA/pH6 has specific cellular actions distinct from HBSS/pH6. *Upper panel* ΔF/F_0_ averaged response of Fluo-4 labeled rCCA perfused first with HBSS/pH6 and then with NMDA/pH6. *Lower panel* ΔF/F_0_ averaged response of Fluo-4 labeled rCCA perfused first with NMDA/pH6 and then with HBSS/pH6. Bars above each curve indicates treatment and its duration.

## RESULTS

### Flux-independent iCa2+ response is elicited mainly by pH6

Given our previous findings describing a flux-independent iCa2+ rise in rCCA in response to 1 mM NMDA at pH≈6 (from now on abbreviated NMDA/pH6) mediated by the NMDAR, we further explored the role of pH in this response. We treated rCCA with 1mM NMDA in HBSS at pH≈7, 6 or 5. It must be noted that at the synaptic cleft, with intense synaptic activity, Glu reaches such concentration and pH falls to values ≈6.4 (12, 15), whereas more acidic pH values (≤5) can be reached under certain pathological conditions (13, 22). As shown in figure 1A (left column), the maximal averaged response was observed with NMDA/pH6 (∫(ΔF/F_0_)=5.49). In contrast, NMDA/pH7 only slowly and slightly increased iCa2+ (∫(ΔF/F_0_)=2.81), whereas NMDA/pH5 also increased iCa2+ (∫(ΔF/F_0_)=3.37), but to a lesser extent than NMDA/pH6. It must be noted that these figures are population averaged responses, but that a diversity of cellular responses was obtained. Please refer to Supplementary Figure 1 where individual cell responses are shown with distribution histograms of population ∫(ΔF/F_0_). In Table I the number of experiments, cells (n) and ∫(ΔF/F_0_)± are shown. We also found that the iCa2+ averaged response to NMDA/pH5 and NMDA/pH6 decreased abruptly after washout, suggesting that the iCa2+ response is ligand binding dependent, in contrast, NMDA/pH7 was not decreased after washout. Considering these results, we further analyzed the role of acidic pH alone on the iCa2+ response treating rCCA with HBSS at pH7, 6 or 5. As observed in figure 1A (right column), HBSS/pH6 elicited a maximal averaged response very similar in magnitude and kinetics to that observed with NMDA/pH6 (∫(ΔF/F_0_)=5.85), although with a marked initial peak that fell gradually after a couple of minutes, response that was not observed with NMDA/pH6, that fell gradually until washout. On the other hand, the iCa2+ averaged responses to HBSS/pH5 and HBSS/pH7 were more modest than those observed in presence of NMDA (∫(ΔF/F_0_)=1.72; ∫(ΔF/F_0_)=0.62, respectively). Indeed, responses to NMDA/pH7 and HBSS/pH7 presented different kinetics, since HBSS/pH7 only slightly increased iCa+ after several seconds of perfusion, but similarly to NMDA/pH7, this response was not reduced after washout. Again, it must be noted here that population averaged responses are shown in these figures, but that a diversity of cellular responses was obtained. Please refer to Supplementary Figure 1 to explore them.

In our previous work we demonstrated that the iCa2+ response of rCCA to NMDA/pH6 is mediated mainly by a flux-independent function of the NMDAR (9). Thus, we tested the role of this receptor to HBSS/pH6 using two different NMDAR competitive inhibitors. As shown in Fig. 1B, and consistently with our previous findings, APV (100μM) and KYNA (20 μM) importantly blocked the averaged response to HBSS/pH6. (∫(ΔF/F_0_)=1.28; ∫(ΔF/F_0_)=1.21, respectively), supporting the role of the NMDAR in rCCA response to acid pH. Again, it must be noted here that population averaged responses are shown in figures, but that a diversity of cellular responses was obtained. Please refer to Supplementary Figure 1 to explore them.

Taken together, these observations demonstrate that the iCa2+ response to NMDA/pH6 or pH5 is mainly elicited by the pH and that only a small component of these responses is related with NMDA itself. In contrast, the iCa2+ averaged response to NMDA/pH7 was clearly different from that elicited by HBSS/pH7. Interestingly, averaged responses at pH7 with or without NMDA were not finished by washout, indicating that different cellular mechanisms regulating iCa2+ are activated at more acidic pH. Also, these findings let us conclude that rCCA sensing of acidic pH6 that elicits iCa2+ rise is in part mediated by the NMDAR, consistently with our previous work. However, since the response was not fully blocked by NMDAR competitive inhibitors, this indicates that other cellular mechanisms must be involved in iCa2+ dynamics after acidic pH exposure.

### HBSS/pH6 and NMDA/pH6 have different cellular effects

Our results described in the previous section demonstrate that NMDA/pH6 or HBSS/pH6 elicited very similar iCa2+ responses and let us conclude that such response is mainly due to pH6. Since our results here and in our previous work also demonstrate that the NMDAR is involved in pH6 sensing by rCCA, we further wanted to explore the possibility that NMDA may have other unappreciated effects, distinct from pH6 treatment. To test this idea, we challenged rCCA with an application of NMDA/pH6 after initial HBSS/pH6 exposure, or the opposite condition: perfusion of HBSS/pH6 after NMDA/pH6 initial exposure. Interestingly, we observed that the second challenge with HBSS/pH6 or NMDA/pH6 elicited distinct responses (Fig. 2). The application of NMDA/pH6 after HBSS/pH6 increased iCa2+ response with similar kinetics to an initial NMDA/pH6 treatment, although somewhat decreased (Fig. 2 upper panel). In clear contrast, when cells were exposed to HBSS/pH6 after initial NMDA/pH6, the iCa2+ response was importantly diminished, and its kinetics was distinct from an initial HBSS/pH6 treatment (Fig. 2 lower panel). These results demonstrate that despite iCa2+ response to HBSS/pH6 and NMDA/pH6 have similar magnitude and they are dependent upon acidic pH, NMDA elicits specific cellular effects that are evident only after a subsequent challenge with pH6.

### mΔψ depletion requires acidic pH and NMDAR flux

Considering our findings here and those previously published, we then explored the role of acid pH in mΔψ depletion. We treated rCCA with HBSS/pH6 (Figure 3A, triangles) but we unexpectedly found no mΔψ depletion as that elicited by NMDA/pH6 (Figure 3A-D, black line). It must be noted here that as in our previous experiments, population averaged responses are shown in Fig. 3, but that a diversity of cellular responses was obtained. Please refer to Supplementary Figure 2 where rasters of individual cell responses are shown along with distribution histograms of the rate between t180/t0 for each cell. Also, in Table II the number of experiments, cells (n) and average ratios ±SEM of t180/t0 are shown. We then treated the cells with NMDA/pH7.3 and we observed that mΔψ was not depleted but instead it slightly increased after several seconds (Fig. 3B, circles). Then we tested the role of the NMDAR on mΔψ depletion using two inhibitors of this receptor, the Glu site competitive inhibitor APV and the irreversible pore blocker MK-801. We found that a low concentration of APV (10 μM) partially blocked the effect of NMDA/pH6 on mΔψ, whereas a higher concentration of APV (100 μM) fully blocked this response, (Fig. 3C, circles=APV 10 μM; squares=APV 100 μM), thus demonstrating that NMDAR activation by NMDA is necessary for mΔψ depletion, as suggested by the lack of effect with HBSS/pH6. Surprisingly, we found that the NMDAR pore blocker MK-801 also inhibited mΔψ depletion elicited by NMDA/pH6 (Fig. 3D, diamonds), indicating that ionic flux through the NMDAR is also necessary. Please refer to Supplementary Figure 2 where individual cell responses are shown.

**Figure 3.**
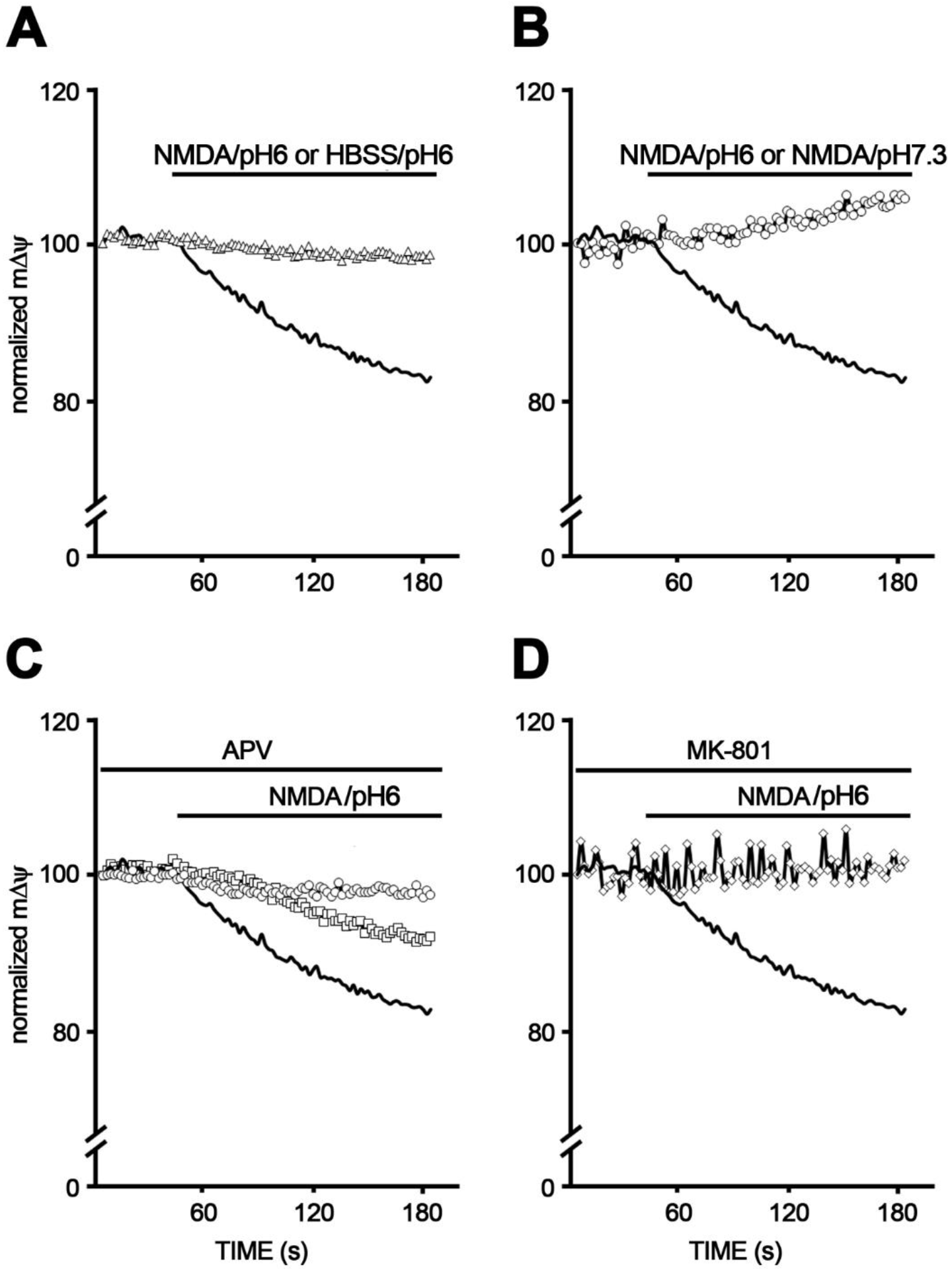
Effect of pH on mΔψ of rCCA and NMDAR role. **A** mΔψ time lapses of rCCA labeled with JC-1 treated with NMDA/pH6 (black line) or HBSS/pH6 (triangles). **B** mΔψ time lapses of rCCA labeled with JC-1 treated with NMDA/pH6 (black line) or NMDA/pH7.3 (circles). **C** mΔψ time lapses of rCCA labeled with JC-1 treated with NMDA/pH6 (black line) with or without APV 10 μM (squares) or 100 μM (circles). **D** mΔψ time lapses of rCCA labeled with JC-1 treated with NMDA/pH6 (black line) with or without MK-801 10 μM (diamonds). All recordings were initiated with 1 min basal recording after which treatments were applied and cells were recorded for two more minutes.

Together, these results demonstrate that both: acidic pH and NMDAR flux-dependent function activated by NMDA are required to deplete mΔψ in rCCA. This clearly contrasts with the flux-independent NMDAR signaling that mediates iCa2+ rise in which mainly acidic pH is responsible. More importantly, our observations so far demonstrate that flux-independent and flux-dependent signaling mechanisms mediated by the NMDAR occur in rCCA and regulate pH sensing and mΔψ depletion, respectively. More importantly, our results indicate that iCa2+ and mΔψ depletion are independent actions mediated by the NMDAR and not dependent, as we concluded in our previous work (Please see discussion). In the following experiments we explored the putative cellular mechanisms that could mediate these functions.

### NMDAR may locate near ER and mitochondria

It is known that subcellular localization of molecules provides specificity to its function, as it has been demonstrated for G protein coupled receptors and channels (23, 24). Therefore, we aimed to identify whether the NMDAR maintains some spatial relationship with the ER and mitochondria in rCCA, given our observations and previous findings from other groups showing ER and mitochondria relationship with the NMDAR in neurons (25–27). For this purpose, we labeled *in vivo* the ER or mitochondria with fixable analogs of MTR or ERT dyes, then fixed the cells and labeled GluN1 with an Ab against its extracellular (EC) domain in non-permeabilized cells, to identify only NMDAR located at the plasma membrane (PM). In these experiments we observed that the ER and mitochondria accumulate mainly in the perinuclear region, as it has been reported previously for the ER (28), although both organelles are also present throughout the cell soma (Fig. 4A and B). In these experiments we also found that GluN1 tends to accumulate in the perinuclear region at the cell membrane. However, given that in this thicker (≥3μm) perinuclear region the ER and mitochondria accumulate and overlap, it is difficult to ascertain whether GluN1-ER or -mitochondria guard a specific spatial relationship. Therefore, we focused our analysis on the distal, flatter and thinnner (≤2 μm) region of the cell soma, where the ER and mitochondria are more disperse, and there is less overlapping by crowding. In this region we observed that some puncta of GluN1 colocalize with the ER or are adjacent to it (yellow arrows Figure 4A inset of Merge image). In addition, GluN1 puncta were also observed in regions where no ER is present. On the other hand, in the distal, flatter and thinner area of the cell we observed that GluN1 is also located near mitochondria, with some GluN1 puncta localized apposed to one tip of mitochondria or surrounding these organelles (yellow arrows Figure 3B inset of Merge image). This observation suggests that in rCCA the NMDAR may locate apposed to mitochondria, as it has been previously demonstrated in neurons, where mitochondria localization presumably facilitates its access to membrane sites of Ca2+ entry (27). Also, and similarly to what we observed with the ER, GluN1 was located in regions of the cell devoid of mitochondria.

**Figure 4.**
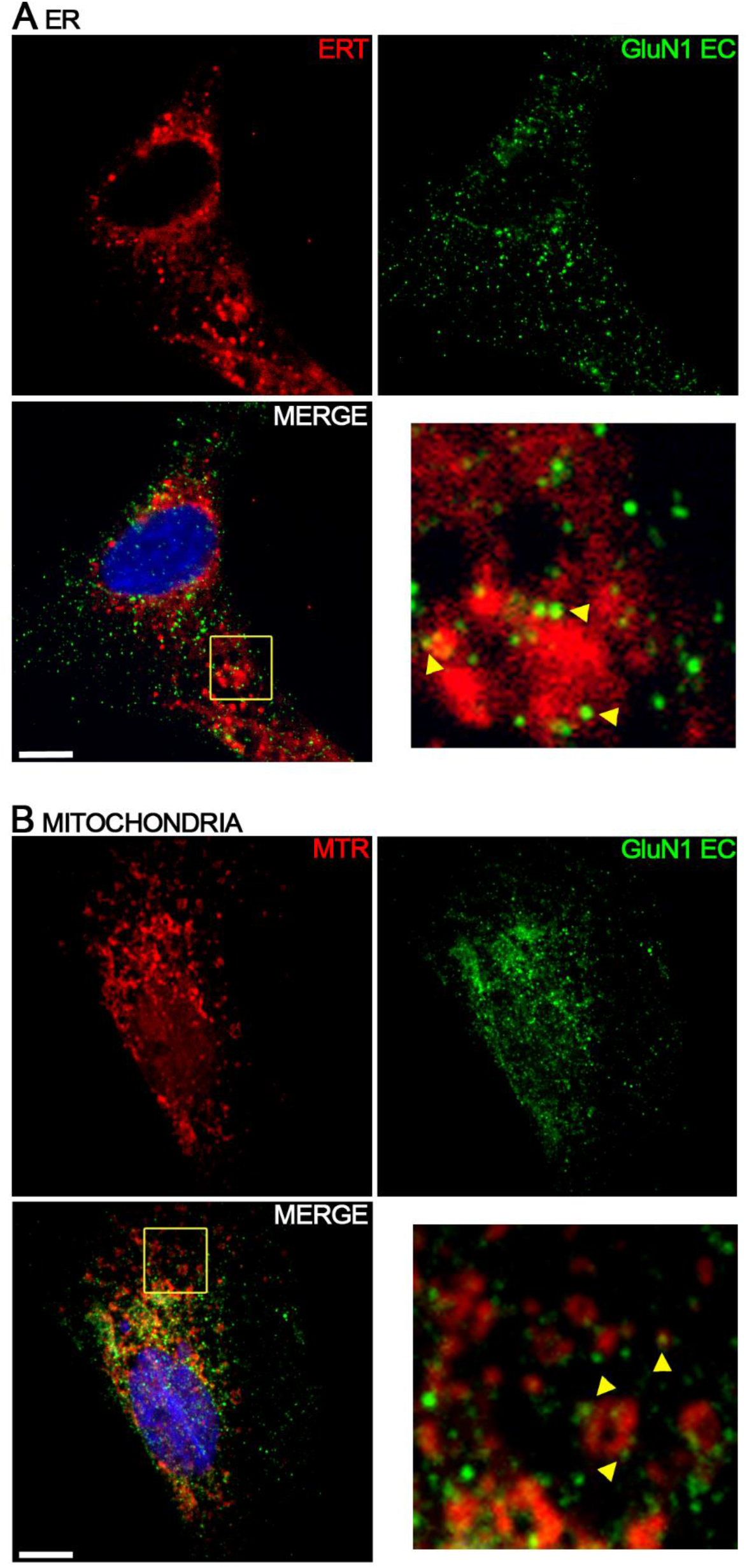
Plasma membrane GluN1 localization relative to ER and mitochondria. **A** EC GluN1 localization relative to ER, in the upper left image it is observed a rCCA labeled with ERT. In the upper right image GluN1 labeling for the same cell is shown. The lower left image shows both channels (Merge) with an inset augmented in the lower right image. Yellow arrows in the inset indicate GluN1 apposed to or surrounding ER. **B** EC GluN1 localization relative to mitochondria, in the upper left image it is observed a rCCA labeled with MTR. In the upper right image GluN1 labeling for the same cell is shown. The lower left image shows both channels (Merge) with an inset augmented in the lower right image. Yellow arrows in the inset indicate GluN1 apposed to or surrounding mitochondria. Reference bar=10μm.

These findings show that some GluN1, and therefore some NMDAR, are located apposed or surrounding the ER and mitochondria in rCCA, making possible that they could be relevant for their regulation.

### pH6 exposure with or without NMDA alters rCCA ultrastructure

Considering our observations and to further gain insight of the cellular mechanisms behind the dual signaling of the NMDAR and the cellular effects of NMDA/pH6 or HBSS/pH6, we analyzed cell ultrastructure by Electron Microscopy (EM). In these experiments we observed that 30 min treatment (time used to exacerbate cellular mechanisms and make them observable by EM) with HBSS/pH6 or NMDA/pH6 changed the density of the cytoplasm, as observed in Figs 5a, c and e, most probably indicating the entry of water and solutes. In addition, we found that both treatments affected the ER, causing swelling or disturbances as it is also observed in Fig. 5c and e, in contrast with the ER from rCCA incubated with HBSS (Fig. 5a), therefore suggesting ER stress. Notably, NMDA/pH6 treatment affected cells at different levels, because in some cells mitochondria were observed within vesicles, hinting autophagia (Supplementary Fig. 3). These observations strongly suggest that pH6 induces the entry of solutes and water into rCCA and generate ER stress that could be the cause of Ca2+ release from the ER, similar to what has been shown in neurons (26). In addition, NMDA caused additional effects consistently with findings described in previous sections.

**Figure 5.**
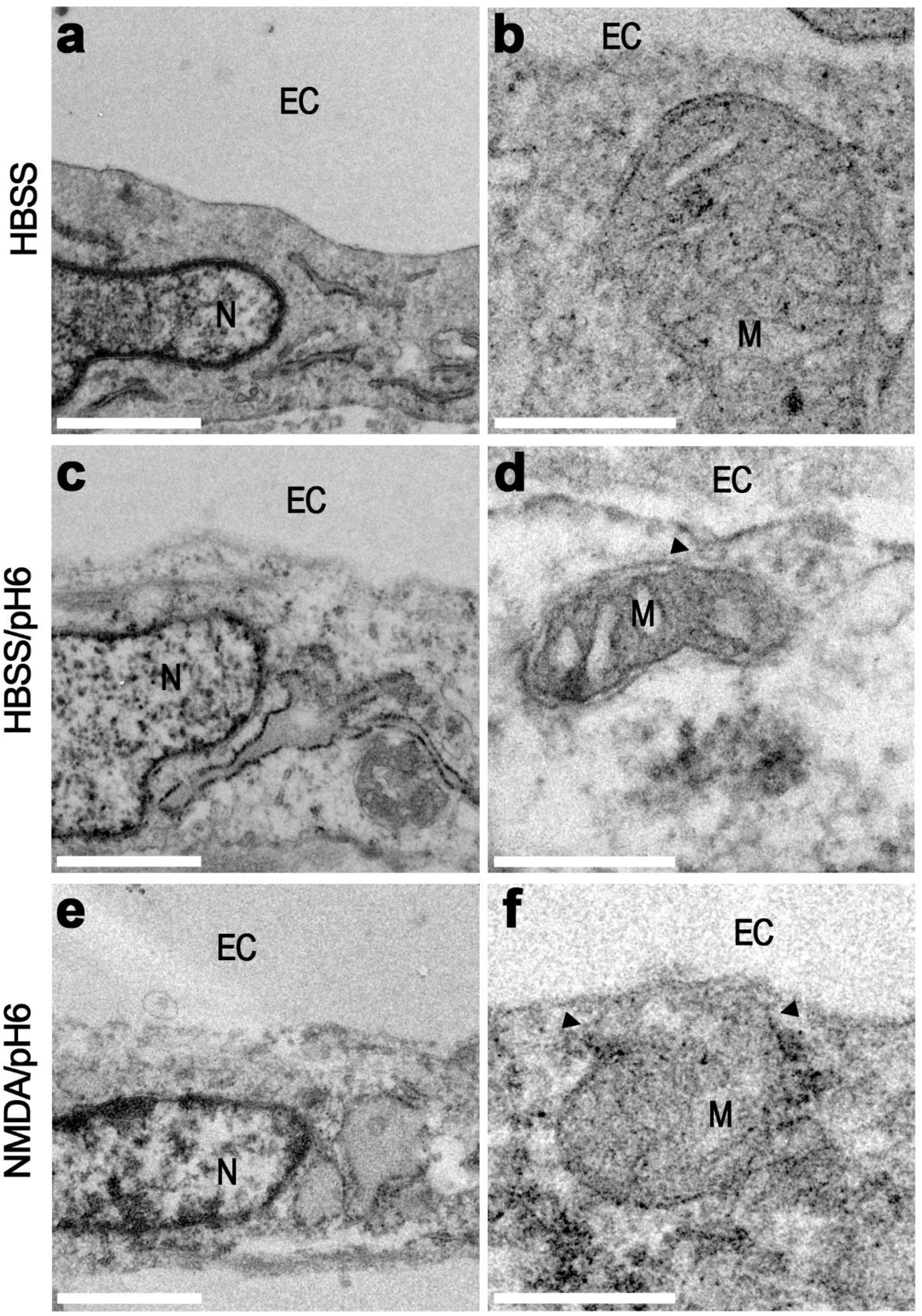
Cellular ultrastructure and mitochondria of rCCA. EM images showing rCCA incubated 30 min with HBSS (**a** and **b**), HBSS/pH6 (**c** and **d**) or NMDA/pH6 (**e** and **f**). In **a, c** and **e** a cell section is shown in which the ER and nucleus are observed. In **b, d** and **f** details of mitochondria and their proximity to the PM are shown. Arrows in **d** and **f** indicate the electrodense areas between PM and mitochondria. N=nucleus; M=mitochondria. Reference bar= 1 μm in **a, c** and **e**; =250 nm in **b, d** and **f**.

### NMDA/pH6 promotes PM-mitochondria bridges (PM-m-br)

EM experiments also showed that mitochondria could be localized very close to the PM (as close as ≈50 nm), as observed in Fig. 5A panel b. This proximity was also observed in cells treated with pH6 with or without NMDA (Figs. 5A panels d and f, respectively). However, we noted that these treatments also generated electrodense areas between the PM and mitochondria (arrowheads in Figs. 5A panels d and f). Furthermore, with NMDA/pH6 treatment, we also observed PM invaginations just above mitochondria (arrow in Fig. 6A panel a), resembling caveola observed in stressed myocytes that presumably mediate caveolin transfer from plasma membrane to mitochondria (18). Moreover, NMDA/pH6 also induced the appearance of large and highly electrodense zones with different size contacting mitochondria and PM (arrowheads Fig. 6A panels b, c, d and e), some of them in direct contact with PM invaginations (arrow Fig. 6A panel b). Strikingly, the larger electrodense zones seem to bridge PM and mitochondria (arrowheads Fig. 6 A panels b and c), because these electrodense areas, beyond contacting these organelles, coincide with darker zones within mitochondria restricted to the contact site (Fig. 6 A panels b, c, d and e). Therefore, we refer here to such structures as PM-mitochondrial bridges (PM-m-br). In contrast, these electrodense zones between PM and mitochondria were only incipient with HBSS/pH6 (arrowhead Fig. 5d), and in our analysis they were never observed as large and electrodense as those observed with NMDA/pH6. Consistently with the observation of PM invaginations elicited by NMDA/pH6, this treatment increased endocytosis of dextran labeled with AF-647 (Fig. 6B upper panel), augmenting the density of vesicles per unit of area that resulted in a larger proportion of the cell covered by these vesicles, that nevertheless presented a smaller average size (Fig. 6B lower panel). Interestingly, with NMDA/pH6 we also observed electrodense zones between mitochondria and ER or nucleus located only a few nm apart (hashtags in Figs. 6A panels d and e, respectively), that in our analysis of control cells and HBSS/pH6 were not observed.

**Figure 6.**
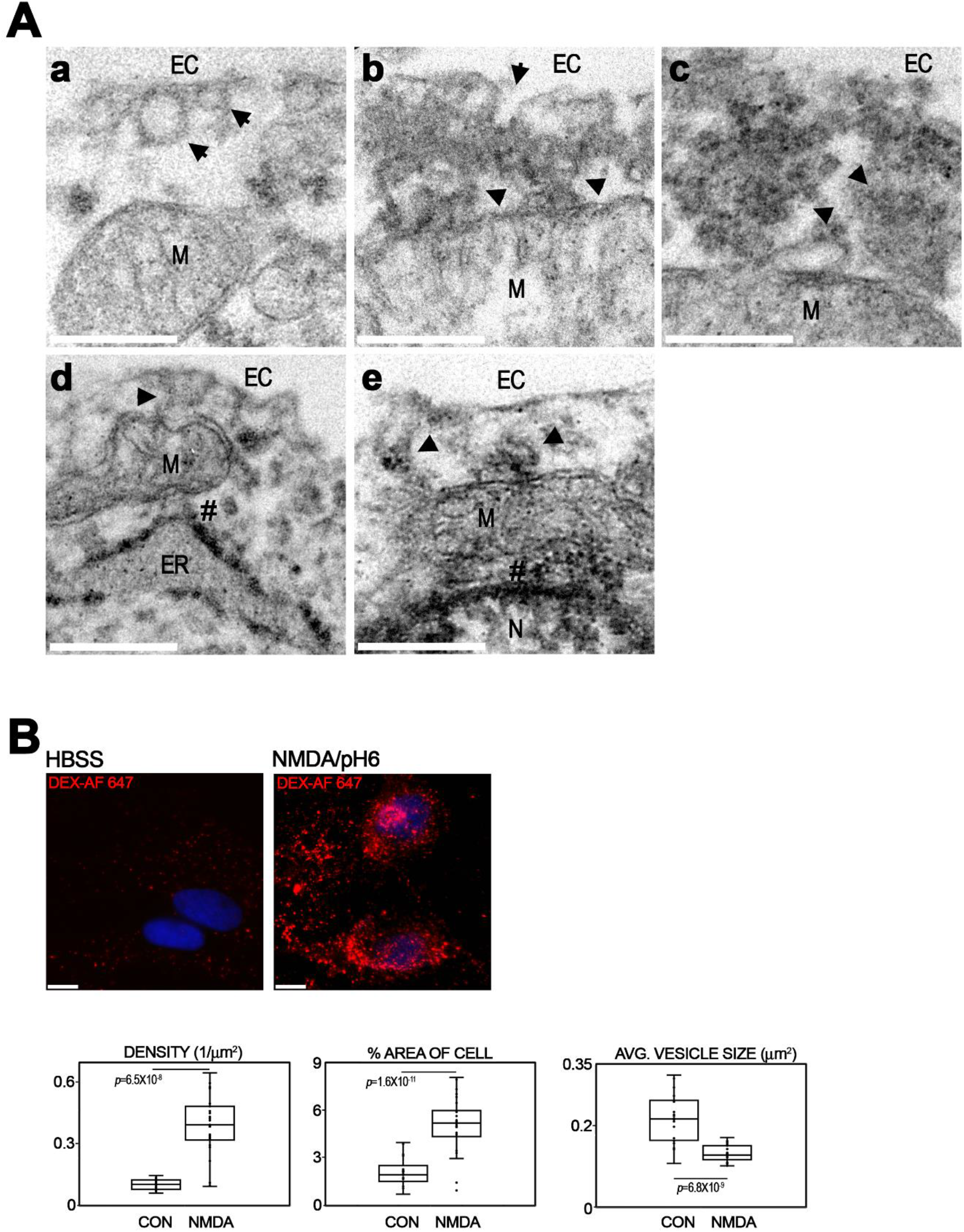
NMDA/pH6 induces vesicles, electrodense structures between PM and mitochondria and endocytosis in rCCA. **A** EM images showing mitochondria of rCCA incubated 30 min with NMDA/pH6. In **a** endocytic vesicles (arrows) can be observed between PM and mitochondria. In **b** and **c** large electrodense structures (arrowheads) or bridges (PM-m-br) are observed between PM and mitochondria. Darker speckled regions are located within mitochondria that coincide with the contacts of these electrodense structures. One invagination (arrow) is observed in **b** in direct contact with these electrodense structures. In **d** and **e** mitochondria located at some nm from ER and nucleus are observed. In between mitochondria and these organelles electrodense zones are observed (hashtags). In in these images electrodense area between PM and mitochondria are also appreciated (arrowheads). N=nucleus; M=mitochondria. Reference bar=250 nm. **B** *upper panel* Representative images of rCCA in which endocytic vesicles filled with AF 647 labeled dextran are shown in control conditions or after NMDA/pH6 treatment. **B** *lower panels* Quantification (average±SEM) of vesicle density per μm^2^; % of cell area covered by the vesicles and average vesicle size (μm^2^). Reference bar=10μm.

Taken together these observations indicate that NMDA/pH6 promotes endocytosis and PM-mitochondria communication through PM-m-br. To our knowledge, the latter has been demonstrated once in other mammalian cells, but has been more studied in yeast in the context of mitochondrial separation between mother and daughter cells (17, 18, 29). Also, NMDA/pH6 increased contacts between mitochondria and other organelles.

Considering previous findings by other groups and mΔψ depletion requirement of NMDAR flux and pH6, our findings suggest the notion that mΔψ depletion could be given by the entry of H+ and/or Ca2+ into mitochondria through these PM-m-br (Please see Discussion). In the following experiments, we further explored PM-mitochondria communication.

### Mass transfer from PM to mitochondria occur in rCCA

As a proof of concept of PM-mitochondria bridging, we tested the transfer of the lipophilic dye FM 4-64FX, a fixable analog of FM 4-64, from PM to mitochondria. This membrane tracer is a lipophilic compound that inserts into the outer leaflet of the PM. After some minutes post-labelling it is internalized attached to the membrane of endocytic vesicles and therefore it can be observed intracellularly. Therefore, for this experiment, rCCA mitochondria were labeled *in vivo* with MTR and afterwards they were incubated for 5 min with FM 4-64FX with or without NMDA/pH6, while the cells were video recorded. In these experiments we observed that after 50 s of incubation, rCCA PM was labeled (Fig. 7 middle column upper and lower panels). Additionally, after 100 s in the presence of NMDA/pH6, rCCA mitochondria were labeled (Fig. 7 lower panel), whereas this was not observed in untreated cells (Fig. 7 upper panel), although different kinetics were observed, thus supporting the notion of PM-mitochondria communication. Importantly, we noticed that not all mitochondria were labeled by FM 4-64, nor that labeling intensity was equal for all mitochondria, thus suggesting that FM 4-64FX transfer to mitochondria is not related with its diffusion through cytoplasm, where presumably free FM 4-64FX is unable to access, as it inserts into the PM. Instead, these experiments support the mass exchange between PM and mitochondria, most probably through the PM-m-br observed in our EM experiments.

**Figure 7.**
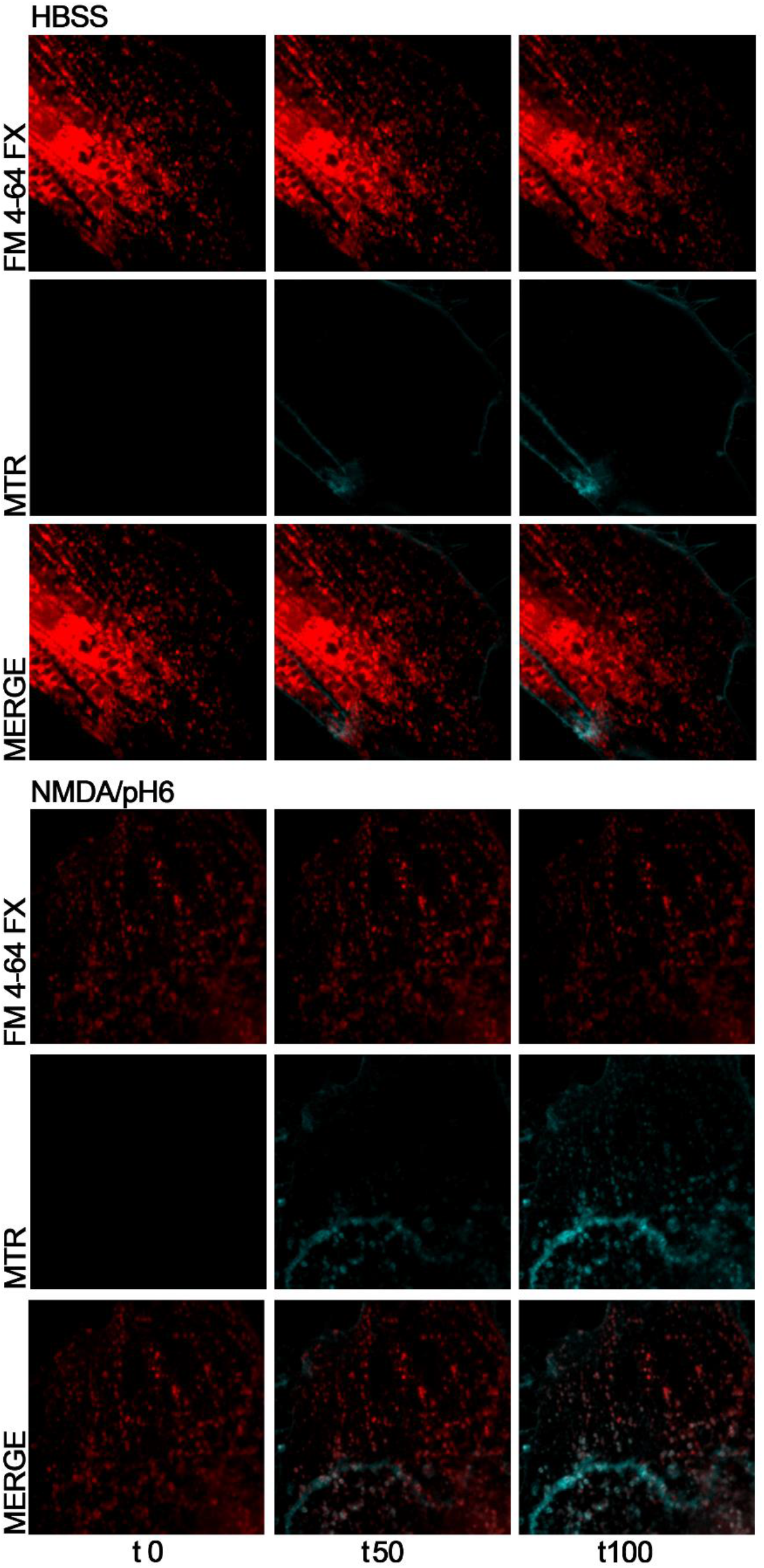
Proof of concept of PM-mitochondria mass transfer. rCCA were labeled with MTR and then incubated 3 min with the lipophilic dye FM 4-64FX while videorecorded. Images of MTR, FM 4-64 and merged images are shown at 0, 50 and 100 s. The upper group shows rCCA incubated with HBSS, and the lower group shows cells incubated with NMDA/pH6. Note that mitochondria are labeled with FM 6-64 after 100 s in the presence of NMDA/pH6, whereas HBSS/pH6 did not induce this phenotype. Reference bar=10μm.

### GluN1 is split apart in rCCA

Our results so far indicated that in rCCA the NMDAR mediates both flux-independent pH sensing and flux-dependent mΔψ depletion. This functional duality could be due to NMDAR with specific subunit composition, given that we have previously shown that all subunits are expressed by rCCA and NMDAR singularities in astrocytes. Alternatively, it is also possible that other mechanisms could be involved. In this regard, it is well known that molecular proteolysis of cell membrane molecules yields IC and EC domains with functions different from the native molecule (30, 31). It has been demonstrated that GluN1 is subject to proteolytic cleavage under different conditions (32–35). Therefore, we explored this possibility analyzing by WB the integrity of GluN1 with two Abs against its IC and EC domains. Importantly, in these experiments we used the same blot to WB both Abs after stripping with a commercial stripping buffer and after signal detection controls were performed. In these experiments both Abs revealed different bands beyond full length GluN1, as observed in Fig. 8A. The Ab against the EC domain of GluN1 detected a conspicuous band around 115 kDa that corresponds to full length GluN1, but also an abundant band around 70 kDa indicated with the letter *a*. On the other hand, the Ab against the IC domain also detected the conspicuous band of 115 kDa corresponding to full length GluN1, but also three clear bands at ≈50 kDa (band *b*), ≈80 kDa (band *c*) and <20 kDa (band *d*). Interestingly, according to the sequence recognition of these Abs (the Ab against the EC domain was generated with the 300 N-terminal aa of GluN1; the Ab against the IC domain was generated with the 50 C-terminal aa of GluN1), the observed MW of these bands is consistent with the theoretical MW of fragments generated after Matrix Metalloprotease or Tissue-type Plasminogen Activator cleavage at sites previously identified in GluN1 (Fig. 8B) (32, 34). In Figure 8C, a schematic drawing of these cleavage sites within GluN1 sequence and the resulting fragments are shown. Band *a* would correspond to the EC fragment generated after cleavage in the stem region before the first transmembrane domain, thus comprising the NTD and LBD domains. Band *b* would correspond to the fragment complementary to band *a*, thus comprising the IC domain, the IC and EC loops and the three transmembrane domains. Band *c* would correspond to full length GluN1 without the fragment generated after the cleavage in the NTD. Finally, band *d* would correspond to the fragment generated after cleaving in the stem region after the EC loop and before the third transmembrane domain, thus comprising mainly the ICD and its transmembrane domain. Intriguingly, the putative complementary fragment to band *d* with an expected theoretical MW of 98.75 kDa is not observed with the Ab against the EC domain. However, it is possible that, given the lack of the ICD and its molecular interactions, this fragment could be readily cleaved in the stem region before the first transmembrane domain, yielding a fragment identical to that in band *a*, that is, this putative fragment would have a short life. Consistently with this possibility, the fragment in band *a* is very abundant, although further experiments are required to test this idea.

**Figure 8.**
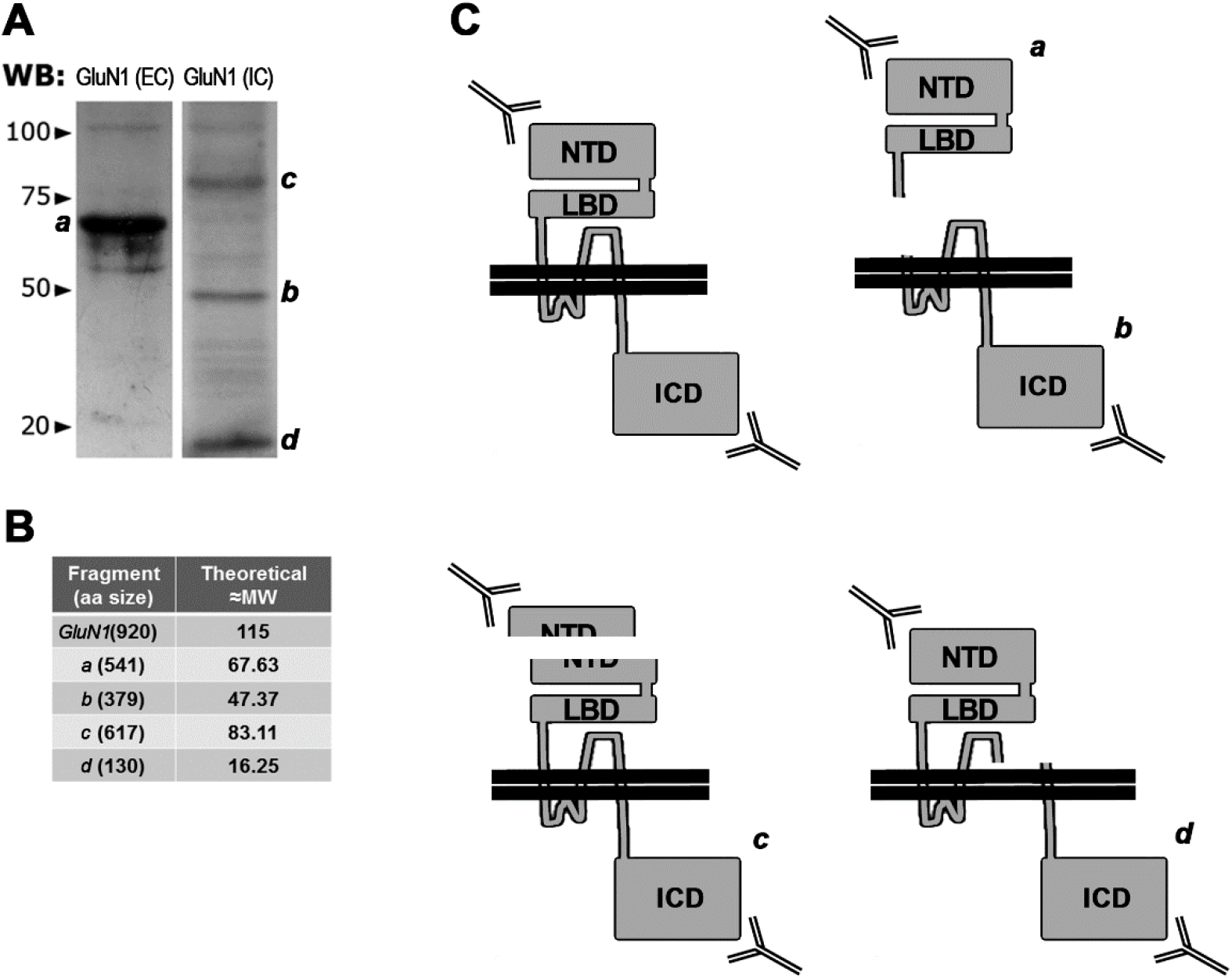
Integrity of GluN1 in rCCA. **A** WB of rCCA lysates probed with the Abs against the EC (*left panel*) and IC (*right panel*) domains of GluN1. The letters indicate GluN1 fragments with different MW identified with these Abs. In the left, arrowheads and numbers indicate the migration of MW markers. **B** Theoretical MW and aa number of full length GluN1 and its smaller fragments identified with letters in A. Theoretical MW was calculated considering cleavage sites previously identified by other groups (please see text). **C** Cartoon representing GluN1, its resulting fragments and domains comprised by each fragment identified with letters in A, with approximate cleavage sites identified by other groups.

These observations suggest that in rCCA GluN1 subunit may undergo proteolysis that splits apart its IC and EC domains, as it has been previously reported in neurons by others in response to NMDA (32). This results in low abundancy of full length GluN1 and the generation of smaller fragments comprising one or more of its domains, thus hinting that different NMDAR populations could be involved in the dual function of this receptor in rCCA.

### GluN1 IC and EC domains are mostly segregated in the cell soma

Since our experiments suggested that in rCCA a large proportion of GluN1 undergoes proteolysis that splits apart its IC and EC domains, we further tested the cellular distribution of GluN1 IC and EC domains with the same Abs used in WB experiments. We found that both Abs showed puncta, with a higher density in the perinuclear region that possibly corresponds to the ER, although the accumulation of NMDAR in the perinuclear region was also observed at PM (see above Fig. 4). We also observed puncta in the cell soma and the cell edge, that most probably correspond to vesiculated and PM NMDAR, respectively (Fig. 9 GluN1 IC and EC images), consistently with the phenotype that we described before (9). However, given the crowding of both labels in the perinuclear region, we centered our analysis in the flat cell soma where we surprisingly observed that both domains are mostly segregated. This is shown in Fig. 9 Merged image, where abundant green or orange dots are observed separately and only scarce yellow dots are found indicating colocalization (yellow arrows, inset of Fig. 9 Merged imaged). These observations confirm that in rCCA GluN1 subunit undergoes some process that splits apart IC and EC domains, as suggested by our WB experiments and as reported by others in response to NMDA (32).

**Figure 9.**
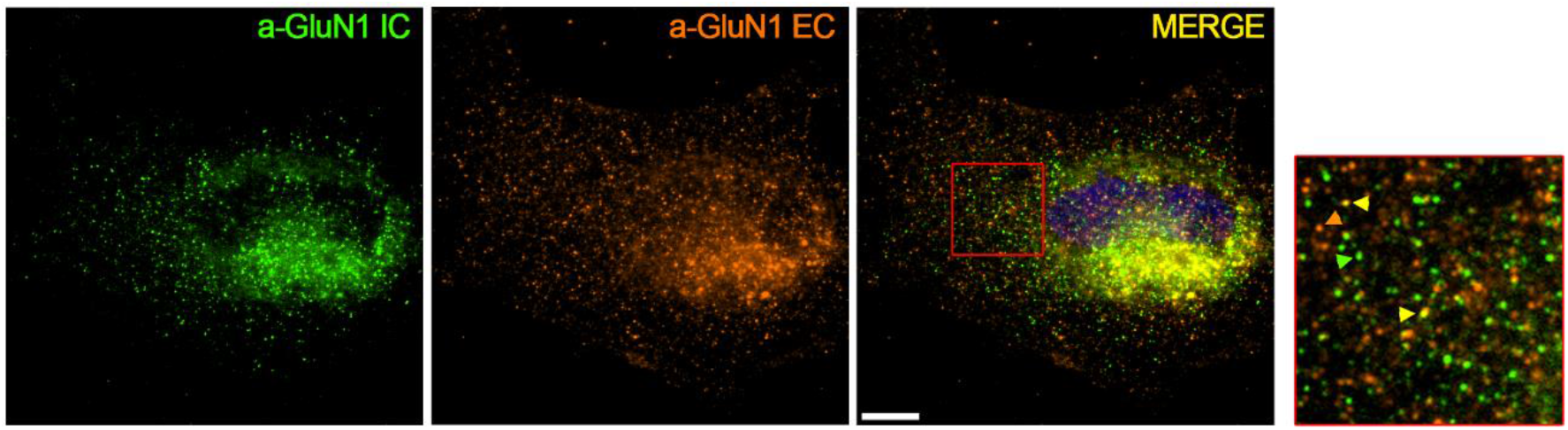
GluN1 IC and EC domain segregation in rCCA. Representative images of rCCA labeled with Abs against GluN1 IC (green channel) and EC domains (orange channel). In the merged imaged both channels are shown, and an inset is indicated. In the enlarged inset, green and orange arrowheads indicate some of the abundant single-color puncta supporting that GluN1 IC and EC domains are split apart. Yellow arrowheads indicate some of the few puncta in which both channels colocalize beyond the perinuclear region. Reference bar=10μm.

Taken together these data confirm that a large amount of GluN1 subunit is cleaved in rCCA, possibly by metalloproteases and/or Tissue-type Plasminogen Activator, given the MW of fragments that fit well with cleavage sites reported previously, although this requires to be demonstrated. These findings further support the possibility that GluN1 processing is involved somehow in the dual signaling properties of the NMDAR in these cells, although this hypothesis still awaits demonstration.

## DISCUSSION

In this work we continued the study of the NMDAR in rCCA, cells that have been demonstrated by others to be devoid of electrophysiological responses mediated by this receptor (8). Nevertheless, several groups have reported cellular functions mediated by the NMDAR (6). We have previously shown that these cells do express NMDAR that mediates iCa2+ rise, similar to findings from other groups, along with mΔψ depletion (9–11). Interestingly, in this work we have found that iCa2+ rise is mainly elicited by acidic pH, rather than NMDA itself. However, and despite iCa2+ responses were very similar between HBSS/pH6 and NMDA/pH6, we unmasked that NMDA also activated cellular mechanisms that are evident only after a subsequent challenge with HBSS/pH6. Such putative mechanisms could include NMDAR endocytosis, consistently with its induction, and/or NMDAR desensitization by posttranslational modifications, since we have shown previously that kinases downregulates its iCa2+ response (9). These findings together with small differences observed by the presence of NMDA at pH 5 and 7, demonstrate that the activation of NMDAR by NMDA exerts specific effects distinct from acidic pH. This was further confirmed by the EM observations that showed clear cellular differences between NMDA/pH6 and HBSS/pH6, including PM-m-br. In addition, we confirmed the participation of the NMDAR in pH sensing because two competitive inhibitors of this receptor reduced importantly the iCa2+ response to HBSS/pH6. This is in good agreement with our previous work, since the iCa2+ response to NMDA/pH6 was also reduced by APV, KYNA, or by a siRNA against GluN1 gene. Nevertheless, it is clear that these inhibitors do not fully eliminate the iCa2+ response, thus suggesting that other mechanisms and/or molecules could participate (see below).

More importantly, in contrast with pH sensing and its iCa2+ response, mΔψ depletion required acid pH and NMDAR flux, thus demonstrating that these functions are mediated by flux-independent and flux-dependent signaling of the NMDAR, respectively. This finding is very relevant because it supports the notion that both functions of the NMDAR may occur concomitantly, as it has been demonstrated in neurons. In this regard, different groups have recently documented that the NMDAR regulates long term depression (LTD) through a flux-independent signaling, whereas long term potentiation (LTP) requires flux-dependent signaling (36, 37)[Reviewed in 5]. These findings open the possibility that similar phenomena of dual signaling may occur in other cells beyond astrocytes and neurons, because it is known that the NMDAR is ubiquitously expressed in different cells and tissues beyond the Central Nervous System (16). Furthermore, since flux-independent signaling mechanisms have been demonstrated for other channels (38, 39), it is feasible that the occurrence of both signaling mechanisms is more common than expected for ionotropic receptors, as some evolutionary traits of the NMDAR also suggest (5). The question then is about the molecular mechanisms that make possible or mediate both signaling strategies in rCCA.

It is well known that the ionotropic function of the NMDAR is regulated by pH (1, 2, 40–43). However, this regulation has the opposite effect of what we observed here, acidic pH inhibits NMDAR flux, and indeed some of the responsible aa have been identified (41). Thus, despite the fact that NMDAR sensing of acidic pH in rCCA represents a novel perspective, the regulation of the NMDAR by pH is not new and therefore it is possible that some of the same aa could be involved. On the other hand, as stated above, since NMDAR inhibitors did not fully eliminate iCa2+ rise, it is possible that molecular interactions of this receptor or other molecules are relevant for this function. In this regard it is known that the NMDAR couples to Ca2+ permeable Acid Sensing Ion Channels (ASICS) (44), and this could be relevant for pH sensing and iCa2+ rise. However, a main direct role of ASICS in iCa2+ rise in response to acidic pH sensing seems unlikely because EC Ca2+ depletion did not eliminate this response, unless flux-independent actions are involved (9). In addition, it has been found that the NMDAR directly interacts with Dopamine receptors, that belong to the G protein coupled receptors family (GPCR) (45). Therefore, it is feasible that different molecular interactions of the NMDAR organized into supramolecular structures at the PM, similar to cytoplasmic signalosomes, could be involved in the rCCA response to acidic pH, however, more experiments are required to answer this question.

The effect of pH in astrocytes has been studied by some groups since some time ago (46). It is known that astrocytes are somewhat resistant to acidic pH (47–49), and this is in good agreement with our observations of rCCA that are capable to survive long periods in acidified media (our unpublished observations). It has been demonstrated that mouse astrocytes of the Retro Trapezoid Nucleus (RTN) respond to pH acidification with iCa2+ increase (50), although the same group found no effect on cortical cultured astrocytes (51), contrasting with our findings. In this regard, it is possible that species-specific effects are responsible for these apparent contradictions, or that particular technical proceedings are behind. A different group found that pH sensing in astrocytes of the RTN is mediated by Kir 4.1 and 5.1 channels (52). With our experiments we cannot rule out that, beyond the NMDAR, these molecules are also involved in rCCA response to acidic pH, however, further experiments are required to test this possibility. In any case, since Kir are K+ channels, their participation would be indirect, unless flux-independent actions of these receptors could be involved. Alternatively, some reports have demonstrated that without ligand, GPCR can be activated by membrane depolarization, thus opening the possibility that this mechanism is involved in iCa2+ rise (53).

One of the most important results of this work is that iCa2+ rise and mΔψ depletion to NMDA/pH6 are mediated by flux-independent and -dependent NMDAR functions, respectively. In our previous work, we erroneously inferred that the mΔψ depletion depended upon iCa2+ increase, because it is well known that Ca2+ uptake by mitochondria through the Mitochondrial Calcium Uniporter (MCU) depletes mΔψ (54). Nevertheless, our findings here clearly demonstrate that this is not the case because iCa2+ rise was mostly related with acidic pH, whereas mΔψ depletion required both: acidic pH and NMDAR flux. Then, iCa2+ rise generated by acidic pH is not sufficient to deplete mΔψ, indicating their independence.

It has been reported that the NMDAR is able to regulate ER and mitochondrial functions (25–27). In particular, the NMDAR regulates ER dynamics in dendritic spines and elicits ER stress. On the other hand, one work reported that in neurons mitochondria are apposed to PM-NMDAR, and this location presumably provides special access of mitochondria to Ca2+ hotspots mediated by the NMDAR. In this regard, it has been shown that molecular location is critical for IC signaling by metabotropic receptors and channels (23, 24), therefore, we investigated whether the NMDAR guards some spatial relationship with these organelles in rCCA. In our experiments we observed that some NMDAR may be located apposed to or surrounding the ER and mitochondria. These observations make possible that the NMDAR is able to regulate the function of these organelles given their proximity. In this regard, we observed by EM that pH6 with or without NMDA generated ER stress. This is consistent with previous findings that reported in mouse cultured cortical astrocytes ER stress in response to acidosis (2-3h), that resulted in cell death after 24-48 h (49). Importantly, it must be noted that in our hands the exposure of rCCA to NMDA/pH6 for up to 30 min did not cause cell death, consistently with the observations by this group who found that 1 h exposure to pH6 did not compromise cell viability. In addition, our EM experiments showed that pH6 with or without NMDA caused a change of cytoplasmic density, perhaps through the entry of solutes and water into astrocytes, that in turn cause ER stress. This observation could be related with the swelling and other cellular transformations that astrocytes experiment during astrogliosis that occurs under different pathological conditions.

EM experiments also showed that some mitochondria in rCCA can be located very close to the PM (at about ≈50 nm). This short distance was also observed in pH6 treated cells with or without NMDA. However, only NMDA/pH6 induced endocytic vesicles between PM and mitochondria, consistently with the increase of endocytosis. These invaginations have the size of and resemble caveolae that have been observed in cardiac myocytes in response to cellular stress by pH, that are related with caveolin transfer from PM to mitochondria, and the increase myocyte adaptation (18). More importantly, HBSS/pH6 and NMDA/pH6 also induced electrodense areas between mitochondria and PM. Strikingly, these areas were only discernible with HBSS/pH6, but they were denser and larger with NMDA/pH6. Moreover, these denser and larger areas observed with NMDA/pH6 are structurally related with PM invaginations, and more importantly, they also coincide with darker electrodense areas within mitochondria, thus we name them here PM-m-br. These observations strongly suggest that PM and mitochondria are capable to communicate and that these structures may mediate mass transfer to mitochondria, as supported by FM 4-64FX labeling of mitochondria, and as Fridolfsson et al. (2012) demonstrated for caveolin in myocytes. Interestingly, functional NMDAR have been reported in isolated mitochondria from neuronal cells (55). Here it is important to note that EM experiments do not exclude that PM-m-br may also occur in control cells, moreover if it is considered glutamate release from rCCA (56, 57). However, it is clear from our analysis that there is an increase of their frequency after NMDA/pH6 treatment.

Further analysis of our data suggests that mΔψ depletion in our experimental conditions could be the result of: a) an increase of H+ within the mitochondrial matrix and/or b) EC Ca2+ entry and uptake by the MCU of mitochondria at the cost of mΔψ(54). The increase of H+ within the mitochondria could be achieved through its uptake from cytoplasm. However, such alternative would require that H+ enter into the cytoplasm first, thus initially increasing mΔψ. Nevertheless, we did not observe such increase with faster frame rate recordings (2.5 Hz; not shown). Therefore, an alternative for the increase of H+ within mitochondrial matrix is the direct entry of H+ from the EC matrix, that could occur through the PM-m-br. On the other hand, mΔψ could be depleted by MCU activity that activates with iCa2+ levels above 10 μM (54), initiated by Ca2+ entry. However, this possibility is unlikely because i) earlier patch clamp studies did not detected Ca2+ flux through the NMDAR in rCCA (8); ii) the rise of iCa2+ elicited by pH6 alone did not deplete mΔψ despite iCa2+ had similar magnitude to NMDA/pH6 and iii) mΔψ depletion requires both NMDAR flux and acid pH. Alternatively, mitochondria could uptake Ca2+ directly from the EC matrix, probably through PM-m-br. In addition, supporting this notion, the establishment of PM-m-br by the astrocyte, that would in principle require cytoskeletal rearrangements considering the force required to form endocytic vesicles, is in the order of minutes, this consistent with the temporal dynamics that require mΔψ depletion (2 min to maximal response) in comparison with Ca2+ rise from the ER (20 sec to peak response). Then, this scenario along with our EM observations and mitochondria staining by FM 4-64 (see below), strongly suggest that mitochondria are capable to uptake H+ and/or Ca2+ from the EC matrix through the structures that we call here PM-m-br, resulting in mΔψ depletion, leading to its stabilization for minutes (9). However, new experiments are required to confirm this idea.

It is possible to think that PM-m-br are the response of astrocytes to NMDA/pH and that they could contain the nanometric tube-like structures that seem to connect mitochondria and caveolae observed by Fridolfsson et al. (2012). Nevertheless, in our conditions in astrocytes, these PM-m-br could represent a more advanced and/or complex state induced by the flux of Ca2+ through the NMDAR. Further considering the density of PM-m-br, it is then possible to conceive that such structures may mediate the transfer not only of PM proteins (18), that most probably include cleaved PM molecules (30), but also of other organic molecules or ions. As a proof of concept of mass transfer from PM to mitochondria, we found that the lipophilic PM dye FM 4-64FX is able to reach shortly mitochondria of rCCA treated with NMDA/pH6, but not in control cells, although different kinetics were observed. In this regard, a difficult but nonetheless possible alternative to this observation is that FM 4-64 labeled mitochondria are located near channels with large pores that enable the diffusion of FM 4-64 into de cytoplasm, and that they are regulated by NMDA/pH6. This would enable direct FM 4-64 capture by mitochondria independently of PM traffic. However, if this were true, we would observe other IC membranal organelles stained with FM 4-64 near these mitochondria, but that is not the case, instead mostly mitochondria are labeled after a couple of minutes.

To our knowledge the communication between PM and mitochondria has only been demonstrated by Fritjolfsson et al. (2012) in mammalian cells, and this is the first time that such structures are observed in astrocytes. Nevertheless, the contact between PM and mitochondria has been investigated in more depth in yeast, in which some molecular players have been identified and it is related with the division of mitochondria between mother and daughter cells (17, 29). In addition, it is possible that the formation of PM-m-br, could help to explain why classical electrophysiology experiments found no response in rCCA to NMDA (8). This is because the movement of ions through the NMDAR could be constrained to the environment of these PM-m-br, that could contain the tube-like structures observed by Fritjolfsson (18). Moreover, the low amount of full length GluN1 that we observed in the cell soma suggest that few complete NMDAR are present, further cooperating to the lack of response in electrophysiological experiments.

From the functional point of view, since our observations strongly suggest that mitochondria employs PM-m-br to directly uptake relevant substrates from the EC space, for instance Ca2+ and H+, these structures would have then a critical role for astrocyte metabolism and survival. Ca2+ is a cofactor of mitochondrial enzymes that promote its metabolism, whereas H+ could be substrate for TCA cycle cofactors to be used in the electron transfer chain, also promoting mitochondrial metabolism and ATP synthesis. This in turn would be crucial to return to homeostasis after stressful stimuli that could lead to astrogliosis. Interestingly, it still remains to be explored if other cells also use PM-m-br to maintain homeostasis, that in the case of transformed cells could be very relevant.

Importantly, in NMDA/pH6 treated rCCA, we also observed electrodense structures between mitochondria and ER or nucleus located at few nm of distance. These observations suggest that in response NMDA/pH6, astrocytes favor physical crosstalk between these organelles in addition to PM-m-br. This could in turn be critical for astrocytes capabilities and responses in different stressful conditions, such as heavy synaptic communication or acidic environment and high EC Glu levels that occur in neuropathological conditions, that may generate astrogliosis. However, new experiments are required to test if these cellular and molecular mechanisms also occur *in vivo* (see below). Again, here we cannot rule out that such crosstalk among organelles occur in untreated cells, we can only assume that their occurrence was promoted. Our findings also support the notion that the communication between organelles occurs more abundantly than conceived (17, 58), and that it occurs in a regulated manner.

Finally, despite the subunits assembled into different NMDAR could provide the basis for its dual signaling described here, it is also known that besides location, posttranslational modifications of PM molecules specify their function. In particular, it is well known that proteolytic processing yields IC and EC molecular fragments with additional functions distinct from those of the native molecule (30, 31). Thus, in an effort to get insight of the putative molecular mechanisms that result in flux-independent and -dependent function of the NMDAR, we explored GluN1 integrity by WB. These experiments showed different MW bands that may correspond to proteolytic sites previously described by others. The split apart of GluN1 was confirmed by the IC distribution of its IC and EC domains. Thus, it is possible that the dual signaling function of the NMDAR is related with its molecular processing by metalloproteases or other proteases present in the culture media. Nevertheless, more experiments are required to clearly establish causality. Interestingly, we identified an abundant molecular fragment of ≈70 Kda that according to epitope recognition by Abs and identified cleavage sites, it may result from proteolysis in the stem region before the first transmembrane domain. This fragment would include the ATD and part of the LBD and seems to remain at the cell membrane perhaps because the molecular interactions mediated by the ATD. This would explain why a soluble GluN1 EC domain was not found in the culture media of neurons, as it occurs with other PM molecules (31, 32). Interestingly, the cleavage at such site is induced by NMDA, and it is known that cultured astrocytes secrete Glu (56, 57), therefore GluN1 cleavage in rCCA could be promoted in an autocrine manner. It still remains to be investigated if other NMDAR subunits are also cleaved. These observations make possible to conceive that the generation of the ≈70 kDa fragment from GluN1 provides the capability to survive the amount of glutamate in the culture media and that secreted by rCCA. This would occur by restricting the amounts of EC Ca2+ that enters the cell through the NMDAR, that when excessive may result into cell death as it occurs in neuronal excitotoxicity. Furthermore, the NMDAR would increase rCCA health by promoting mitochondrial metabolism through PM-m-br direct uptake of EC Ca2+ and/or H+. Evidently, this hypothesis requires further testing, but our observations strongly support it, and if true this novel mechanism observed in astrocytes, and previously in myocytes, could turn out to be crucial for astrocyt’s biology and thus, for neuronal health.

A very relevant question that arises is whether these phenomena occur in actual tissue astrocytes and not only in cultured astrocytes. Despite more research is needed to answer this question, historically we know that many molecular mechanisms are conserved in cultured cells, and cultured astrocytes have demonstrated their utility to study different cellular mechanisms (59). Indeed, in cell biology the exacerbation of cellular mechanisms is a common strategy that facilitates or makes possible their study, such as for instance the use of plasmid transfection to overexpress proteins. These strategies have extensively demonstrated to be useful approaches to elucidate cellular mechanisms common to many molecules (31). In this regard, it is very important to note that the exposure of rCCA to high amounts of NMDA and H+ could mimic specific physiological or pathological conditions. Thus, it cannot be ruled out *a priori* that the cellular mechanisms described in this work could function at small domains of astrocytes *in vivo*, mainly because our observations here are the result of endogenous molecules at work, without molecular overexpression. This leads to speculate which could be these putative astrocyte domains, if they may occur near the synapse, at extra-synaptic sites, near the vasculature or elsewhere. In this regard, the location of Ca2+ stores and InsP3 diffusion in the Peri Synaptic Astrocyte Projections becomes relevant (60, 61)

## CONCLUSIONS

Data presented here demonstrate that the NMDAR in rCCA is able to signal through flux-independent and -dependent manner that regulate Ca2+ release from the ER and mΔψ depletion, respectively. Ca2+ release is mainly due to acidic pH and could be mediated by ER stress, although our data indicates NMDA specific actions, as well as that other molecular entities and cellular mechanisms could be involved beyond the NMDAR. On the other hand, one of the main findings of this work, consistent with previous findings from other groups, is the strong suggestion that mΔψ depletion is mediated by PM-mitochondria communication through the structures referred here as PM-mitochondria bridges (PM-m-br) The generation of PM-m-br require NMDAR ionotropic function and acidic pH, similar to mΔψ depletion, and could mediate Ca2+ and H+ direct access to mitochondria. Given the metabolic capabilities and resilience of astrocytes to stressful conditions, and neuron-astrocyte interactions, PM-m-br could be of great relevance for the neurodegenerative diseases or other pathologies characterized by neuronal death. Moreover, since PM-mitochondria communication has been reported previously in myocytes, it is possible that PM-m-br could be a general cellular mechanism relevant for other cell types and conditions, such as cancer cells. Finally, we found evidence suggesting that GluN1 subunit, the *sine qua non* subunit for all NMDAR, is cleaved, possibly by metalloproteases and/or other Tissue-type Plasminogen Activator. However, new experiments are required to elucidate the functional relevance of this finding for NMDAR dual signaling described here. In addition, more research is needed to study the molecular players that mediate PM-m-br, GluN1 cleavage and their occurrence *in vivo*. This in turn could open new perspectives to tackle brain pathologies in which astrocytes are involved. All in all, our findings open several new questions that require further research to be answered, that could lead to new perspectives of cell biology and some pathologies.

## ACKNOWLEDGMENTS

PMOB deeply thanks Dr. AHC the opportunity to continue his scientific career at his lab at Instituto de Fisiología Celular, UNAM, given the difficulty and poor working conditions at INNN. Also, the authors wish to specially thank M. in Sc. Rodolfo Paredes Diaz for his technical expertise for sampling processing for EM and Dr. Abraham Rosas Arellano for his insightful comments on EM, both from the imagery hub at Instituto de Fisiología Celular, UNAM. The authors also wish to thank Dr. Ruth Rincón Heredia and Dr. Fernando García from the imagery hub from Instituto de Fisiología Celular, UNAM. The authors also wish to specially thank MVZ Cesar Augusto Rodríguez Balderas from the animal house at INNN. The authors declare no conflict of interest. This work was supported by Grant 132706 from CONACyT for Basic Science granted to PMOB in 2010. PMOB wishes to deeply thank his family that has supported him despite lack of conditions to do science at INNN.

---

**Supplementary Figure 1**. Raster plots and ∫(ΔF/F0) distribution histograms for individual cells from experiments of iCa2+.

**Supplementary Figure 2**. Raster plots and t180/t0 rates distribution histograms for individual cells from experiments of mΔψ.

**Supplementary Figure 3**. EM microphotograph of a rCCA treated with NMDA/pH6 in which vesicles containing mitochondria are observed. Bar= 1μm.

