## Supplementary material for "Dual NMDAR signaling in astrocytes: flux-independent pH sensor & flux-dependent mitochondrial regulator through membrane-mitochondria communication": Suppl. Figures 1-3

SUPPLEMENTARY FIGURE 1

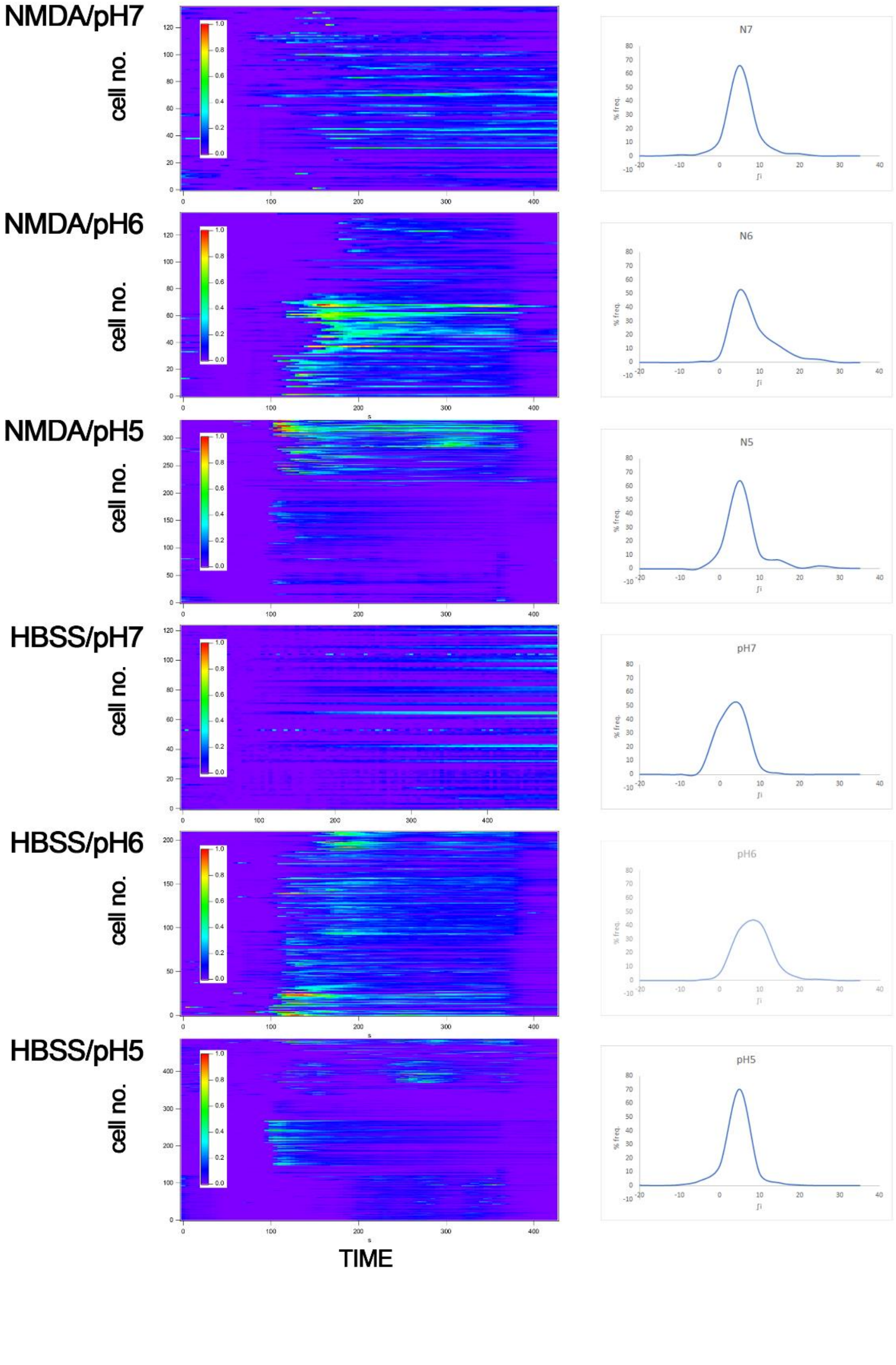

SUPPLEMENTARY FIGURE 1

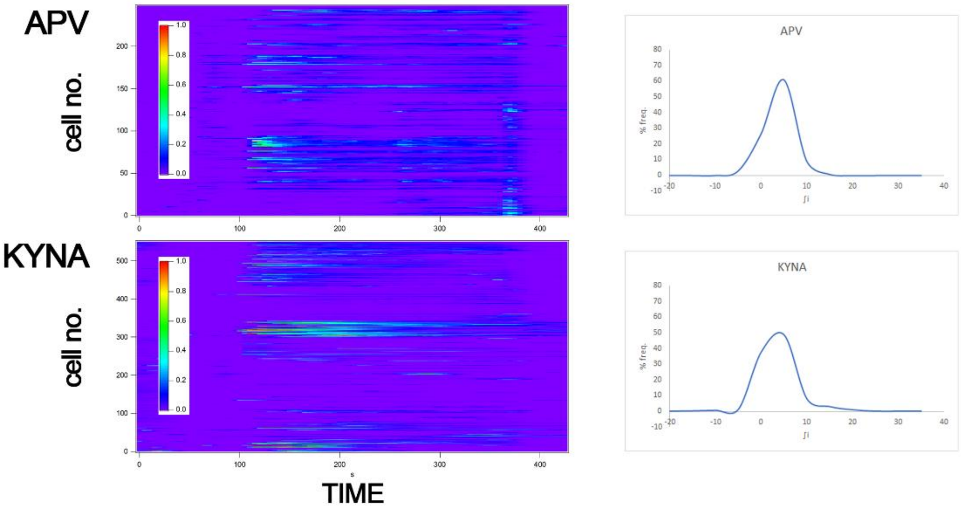

SUPPLEMENTARY FIGURE 2

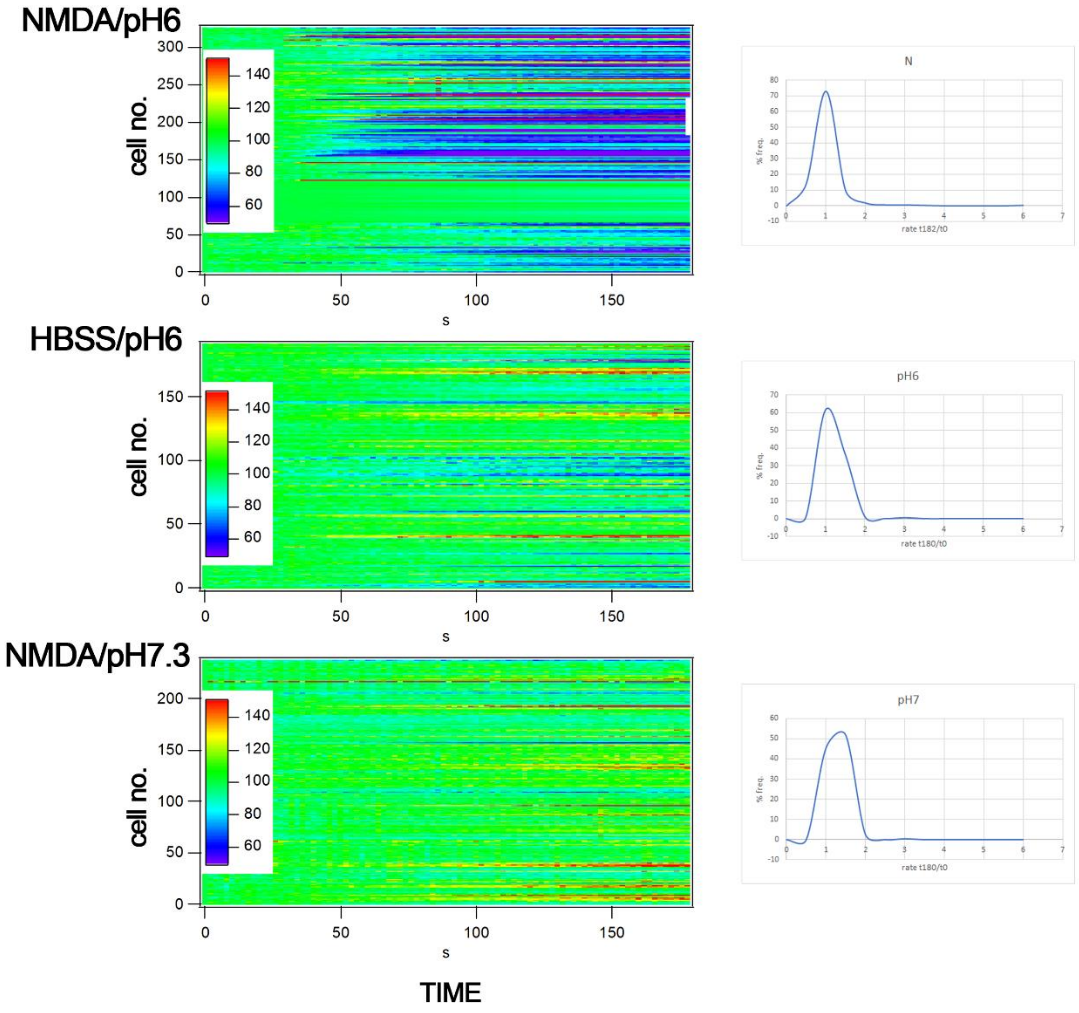

SUPPLEMENTARY FIGURE 2

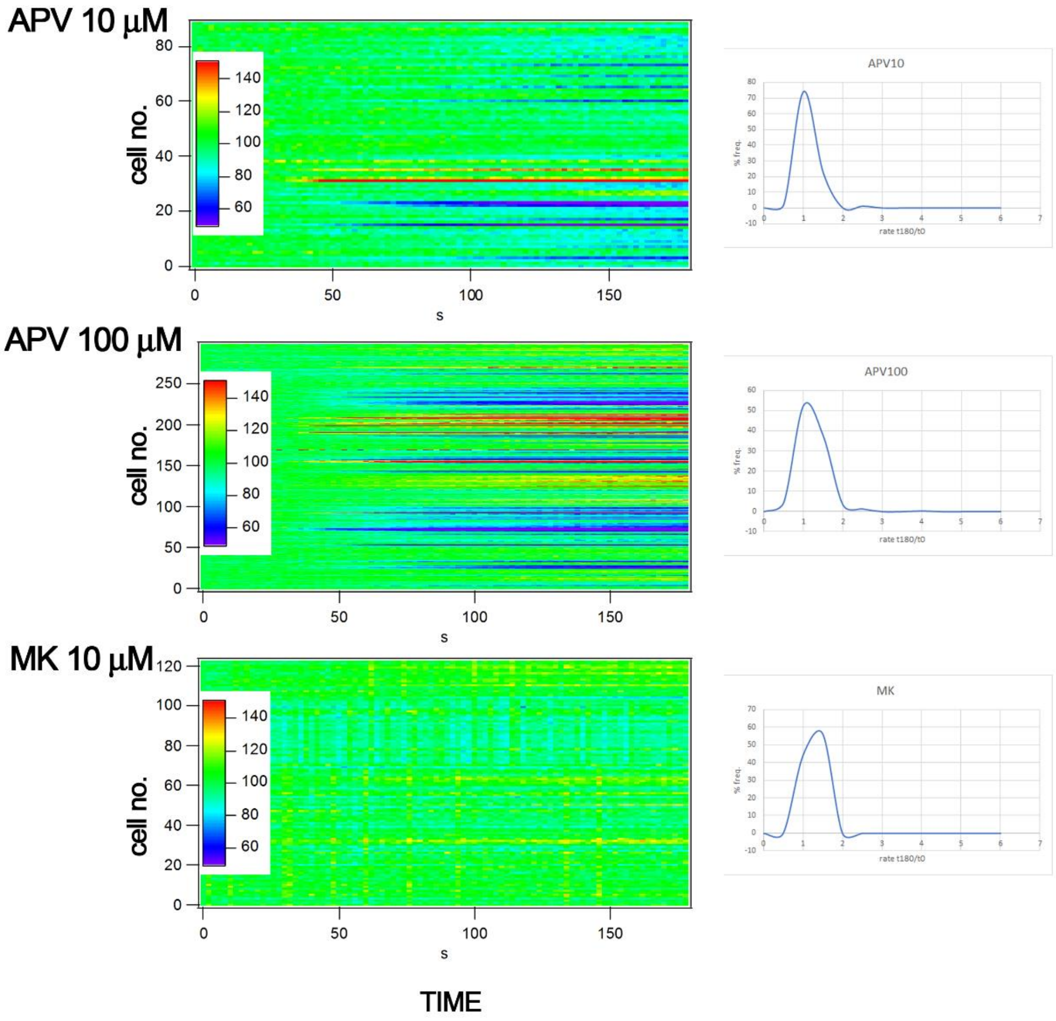

SUPPLEMENTARY FIGURE 3

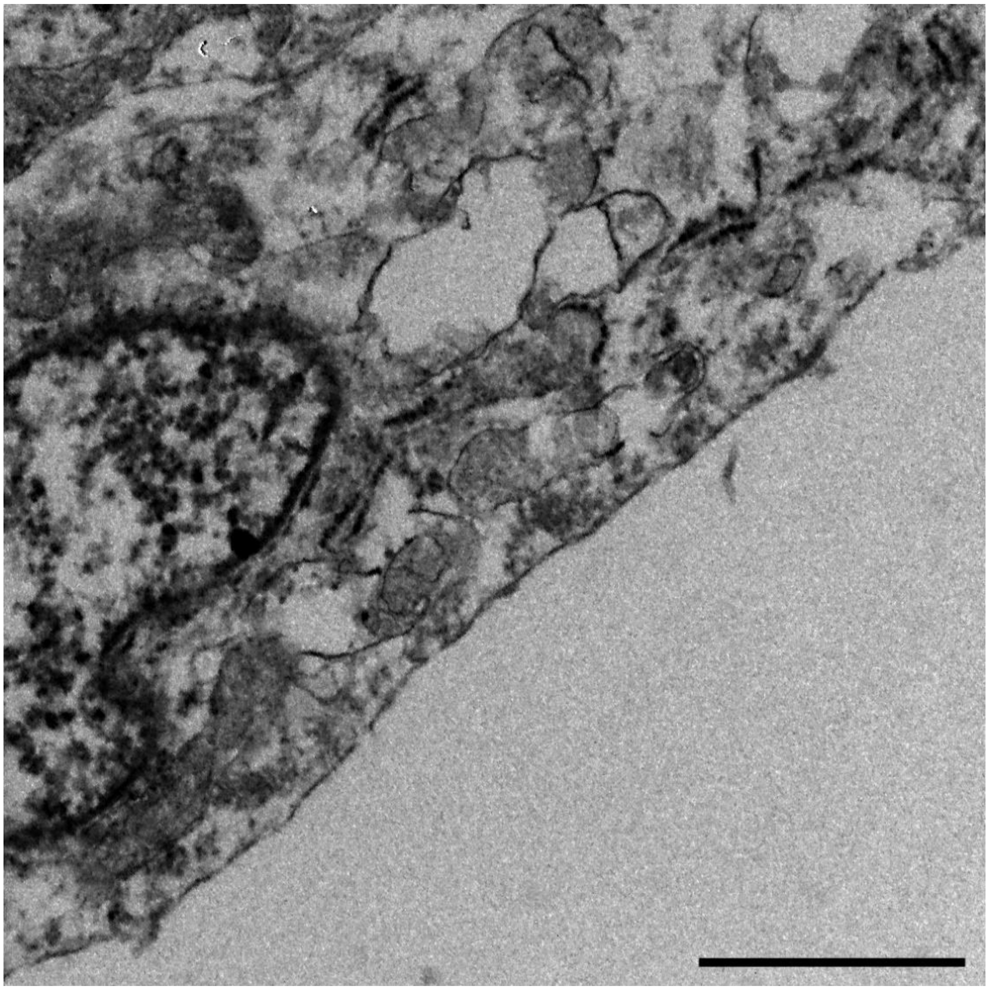
